# Bridging Neurons to Behavior: a Generative Neural Engine Reveals Violations of Independence in the Stop Task

**DOI:** 10.64898/2026.05.09.723954

**Authors:** Alberto Tubito, Andrea Ciardiello, Cristiano Capone, Giampiero Bardella, Pierpaolo Pani, Stefano Ferraina, Guido Gigante

## Abstract

The brain produces robust, low-dimensional behaviour from variable, high-dimensional activity. The dominant account of action control, the Independent Race Model (IRM), explains stopping as a “Go” and a “Stop” process racing independently, and predicts behaviour accurately. But independence is a behavioural assumption never confronted with neural dynamics, and the same premotor neurons drive both processes. We therefore trained a generative Deep Markov Model on premotor activity from two macaques; trained on neural data alone, it emergently reproduces the full reaction-time distribution, validating it as a proxy for the neural hardware. Used for in silico experiments, this engine reveals systematic violations of both of the IRM’s independence axioms, with violations emerging from the geometry of a single shared manifold. Apparent independence was never real: an artifact of observing behaviour without seeing the manifold that produces it.

## 1 Introduction

Complex biological systems present a fundamental paradox: how does stable, robust function—like a thought, a heartbeat, or a motor command—arise from a system built of chaotic, noisy, and variable low-level components (Faisal et al., 2008; Kitano, 2004)? A modern synthesis proposes that high-level function imposes “downward causation” (Noble, 2012) to constrain and compress millions of degrees of freedom into a compact “interface” with the environment (Branchi, 2025).

A useful framework for understanding this compression comes from the physics of complex systems: in many biological models, only a few “stiff” dimensions tightly govern function, while the vast majority of dimensions are “sloppy”—the system’s behaviour is robust to their variation (Gutenkunst et al., 2007; Machta et al., 2013; Transtrum and Qiu, 2014). Identifying these stiff dimensions is the key to understanding how function emerges.

In motor control, this stiff/sloppy structure is directly visible, notably in premotor and motor cortices. Although action stopping engages a distributed network spanning basal ganglia and prefrontal cortex, dorsal premotor cortex (PMd) is positioned at its output interface, such that much of the wider network’s behaviourally decisive computation may be funnelled through local PMd dynamics. Population recordings from PMd reveal that neural activity is, in fact, confined to a low-dimensional manifold (Gallego et al., 2017; Sadtler et al., 2014): movement initiation corresponds to a neural trajectory escaping a preparatory subspace (Churchland et al., 2012; Genkin et al., 2025), while successful inhibition corresponds to the arrest and reversal of that trajectory (Pani et al., 2022; Giarrocco et al., 2021; Bardella et al., 2024).

These descriptions, however, rely on dimensional reduction methods such as PCA (Jolliffe, 2002), which compress the data but do not learn its temporal rules. We can describe a trajectory on a latent manifold, but we cannot simulate in this compressed space and produce behaviours—like reaction times, for example. The generative link between neuronal manifold and behaviour has been missing.

This gap maps onto a classic tension in neuroscience. The field’s dominant behavioural theory of action stopping—the Independent Race Model (IRM) (Verbruggen and Logan, 2009)—posits that a “Go” process and a “Stop” process race independently to a threshold. The IRM has been tremendously useful for estimating quantities such as the stop-signal reaction time (SSRT), but its core assumption—that the two processes do not interact—is difficult to reconcile with what we know about PMd: the same neurons participate in both initiation and inhibition within a single, heavily recurrent network (Boucher et al., 2007; Pani et al., 2022). The tension is one of levels: the IRM is itself a low-dimensional model, but of behaviour—it says nothing about the high-dimensional neural dynamics from which behaviour emerges. To adjudicate it, one needs a low-dimensional model of the neural side that is nonetheless faithful to behaviour; such a model could test the IRM’s independence assumptions against the dynamics that generate behaviour, not merely against behaviour itself.

This level mismatch is not merely conceptual—it shows up in the data. Indeed, converging evidence from partial-response electromyography (Fiori et al., 2025; Raud et al., 2022), kinematic tracking (Hannah et al., 2023), and systematic behavioural analyses (Bissett et al., 2021b) has increasingly pointed to departures from independence, as posited by the IRM, at the single-trial level, while copula-based theoretical work has shown that certain forms of dependence are compatible with existing mean SSRT estimates (Colonius and Diederich, 2018; Colonius et al., 2024). In particular, Bissett et al. (2021b) demonstrated that stop-failure RTs deviate systematically from the no-stop RT distribution in an SSD-dependent manner—positive at short SSDs and negative at long SSDs—a pattern that challenges the model’s assumption of context independence. What has been missing is a generative “Algorithm” in the sense of Marr (Marr and Poggio, 1976), bridging the Computational and Implementation levels—precisely the low-dimensional neural model called for above, now made generative.

Here we ask: can a compact dynamical system, learned entirely from neural activity, serve as a mechanistic probe of the independence the IRM assumes? Two properties are essential for such a system to constitute a genuine test. First, it must be low-dimensional: the dynamics should live in a small number of variables, not merely be projected onto them. Second, it must be Markov: the current state alone must suffice to predict the future, without requiring a long memory of past states. A system that is low-dimensional but non-Markov is not truly compact—its effective dynamical dimensionality scales with the memory depth. If a model satisfying both requirements, trained exclusively on neural activity, can achieve zero-shot prediction of the animal’s behaviour, it becomes a validated proxy for the neural hardware—and we can use it to ask whether the independence posited by the IRM is realized, or not, at the level of the underlying dynamics.

We show that a 3-dimensional Deep Markov Model (DMM) (Krishnan et al., 2017)—a probabilistic framework that learns the hidden dynamical rules of a system from time-series data—serves as this test when trained on PMd multi-unit activity. This engine recreates the monkey’s complete RT distribution from its intrinsic dynamics alone, and—used as a ‘virtual brain’ for in silico experiments—reveals systematic violations of both independence axioms of the IRM: context independence, through a simulated stop-failure analysis consistent with the Pause-then-Cancel framework (Schmidt and Berke, 2017; Diesburg and Wessel, 2021), and stochastic independence, through a per-trial analysis revealing correlated Go and Stop speeds. Both violations emerge from the geometry of the shared neural manifold, even as the IRM continues to provide an accurate and useful account of behaviour at the aggregate level.

## 2 Results

To reverse-engineer the generative algorithm driving motor control, we analyzed the multi-unit activity (MUA) signal derived from broadband neural recordings in the dorsal premotor cortex (PMd) of two macaque monkeys performing a countermanding reaching task. Our analysis proceeds in five main steps. First, we establish the behavioural task framework. Second, we benchmark our nonlinear DMM against descriptive linear models (PCA), demonstrating its superior “dynamical predictability.” Third, we evaluate models of varying dimensions to establish 3D as the minimal dimensionality required for functional completeness. Fourth, we show this 3D engine emergently recreates the full behavioural RT distribution, validating it as a sufficient generative model of movement initiation. Finally, we utilize this validated “virtual brain” as an in silico simulation environment first to mechanistically test the assumptions underlying the Independent Race Model (IRM), and then to perform causal neural perturbations.

### 2.1 Experimental Framework: The Countermanding Reaching Task

The foundation of our analysis is a dataset of multi-unit activity (MUA) derived from broadband neural recordings in the left PMd of two rhesus macaques using a 96-channel Utah array. The animals were trained to perform a countermanding reaching task, a paradigmatic framework for investigating the competition between movement initiation and inhibition (Fig. 1).

**Figure 1.**
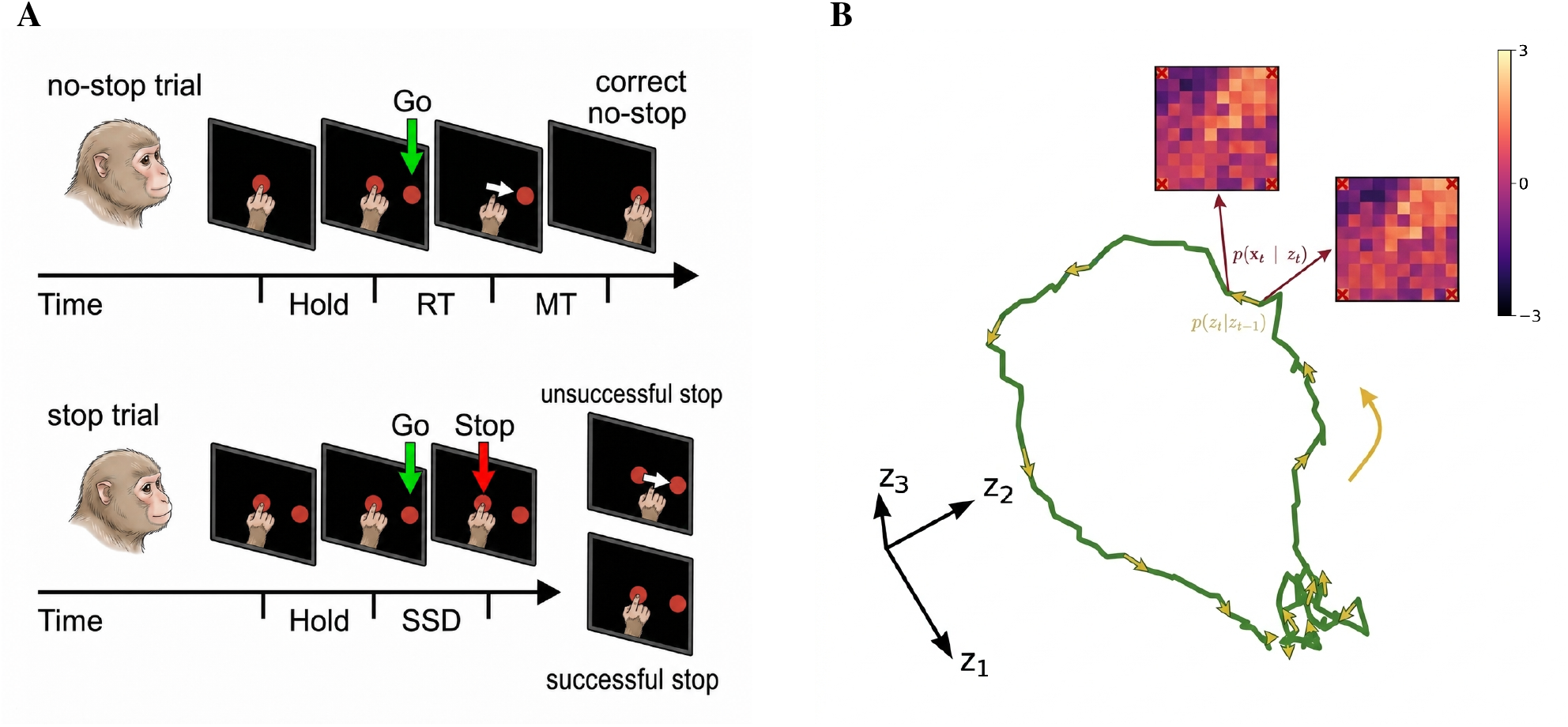
From behavioural paradigm to generative neural engine. **(A)** The countermanding reaching task. Monkeys initiated each trial by holding a central target. In no-stop trials, the disappearance of the central target and the simultaneous appearance of a peripheral target instructed the monkey to execute a reach, to either left or right. Reaction time (RT) was defined as the interval between the Go cue and movement onset. In Stop trials, the central target reappeared after a variable Stop-Signal Delay (SSD), instructing the monkey to inhibit the planned movement. Trials were classified as successful Stops if the movement was cancelled, or unsuccessful Stops if the movement was executed despite the Stop signal. **(B)** The learned generative engine in latent space. Example 3D latent trajectory from a representative no-stop trial. The Deep Markov Model (DMM) learns a stochastic transition model (yellow arrows) that defines a low-dimensional dynamical flow field governing the temporal evolution of latent states. A separate decoding model (red arrows) maps latent states back to instantaneous multi-unit activity (MUA); the two example MUA maps shown are model-decoded reconstructions, arranged according to the spatial layout of the electrode array, with color encoding z-scored MUA (colorbar, range [− 3, 3]). Corner positions marked with a red cross correspond to array sites without a recording electrode.

Each trial began with the monkey holding a central red target. In the majority of trials (no-stop or Go trials), the central target disappeared while a peripheral target appeared simultaneously to the left or right. This event served as the “Go” cue, instructing the monkey to initiate a reach toward the peripheral target. The time elapsed between the Go cue and the onset of movement, either left or right, is defined as the reaction time (*RT*).

In a subset of trials (20–30%), termed Stop trials, the central target reappeared after a variable Stop-Signal Delay (SSD), instructing the animal to cancel the planned movement. Stop trials yielded two distinct outcomes: “Successful Stop” trials, where the monkey inhibited the reach, and “Unsuccessful Stop” trials, where the movement was executed despite the Stop signal.

### 2.2 Three Markov dimensions are necessary and sufficient to capture PMd motor dynamics

To capture the system’s generative dynamics, we trained a Deep Markov Model (DMM) on the multi-unit activity.

Fig. 1B visualizes the core of this approach using a representative single-trial rollout. The DMM generates a low-dimensional latent trajectory (green line) that is not merely a descriptive path, but the result of an active dynamical process akin to an engine of movement (Churchland et al., 2012): it evolves stochastically through a learned low-dimensional force field (yellow arrows), driven by intrinsic noise. This internal engine drives the system forward in time, while a separate decoding step (red arrows) maps these latent states back to the instantaneous, noisy neural activity.

We then tested if this nonlinear stochastic model could capture the system’s generative dynamics more effectively than a standard, descriptive linear model (PCA). We found that while both models can describe neural variance, only the DMM captures the system’s dynamics.

We first compared the ability of a DMM and a PCA model, each with a 3-dimensional latent space, to capture the neural dynamics (Fig. 2); we refer to these as the 3D DMM and 3D PCA, respectively. While PCA is a standard tool for reducing the dimensionality of neural data, it is inherently limited to “compressing” spatial information—providing a snapshot of the manifold—without learning the rules of its evolution. The DMM, conversely, learns the vector field that drives the system. This distinction is quantified in Fig. 2B. While both models reconstruct the static neural activity (*x*_1:*T*_) with similar accuracy (Fig. 2A), their ability to capture the temporal evolution of the system is vastly different (Fig. 2B). When we attempted to predict the next latent state *z*_*t*+1_ from the current history *z*_1:*t*_ (see Sec. 4.4), the PCA-derived space offered limited predictive power 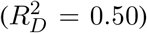. Here, 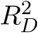 is a multidimensional generalization of the standard *R*^2^ for *D >* 1; it measures the explained variance of the system’s temporal evolution by quantifying how much of the future state prediction error is accounted for relative to the natural noise structure (covariance) of the system’s innovations (Δ*z*) (see Methods Sec. 4.4 for details). This indicates that PCA emphasizes instantaneous spatial patterns over their development over time. The DMM latent space, explicitly trained to implement a dynamics, achieved high dynamical predictability 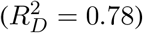 , confirming it represents not just a descriptive subspace, but the system’s generative engine. This result is further strengthened (inset of Fig. 2B) when *z*_*t*+1_ is predicted using only the immediate previous step *z*_*t*_; under these conditions, DMM 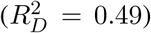 dramatically outperforms PCA 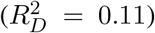. This result is key to our central claim. The one-step (Markov) predictor tests whether the current latent state, by itself, determines the system’s next step—the hallmark of a genuinely low-dimensional dynamical system. The DMM’s strong one-step predictability confirms that its 3 latent dimensions constitute a true 3-dimensional dynamics, not merely a 3-dimensional projection of a higher-dimensional process. PCA’s near-zero one-step predictability, by contrast, indicates that its 3 components, while capturing spatial variance, do not form a self-contained dynamical system.

**Figure 2.**
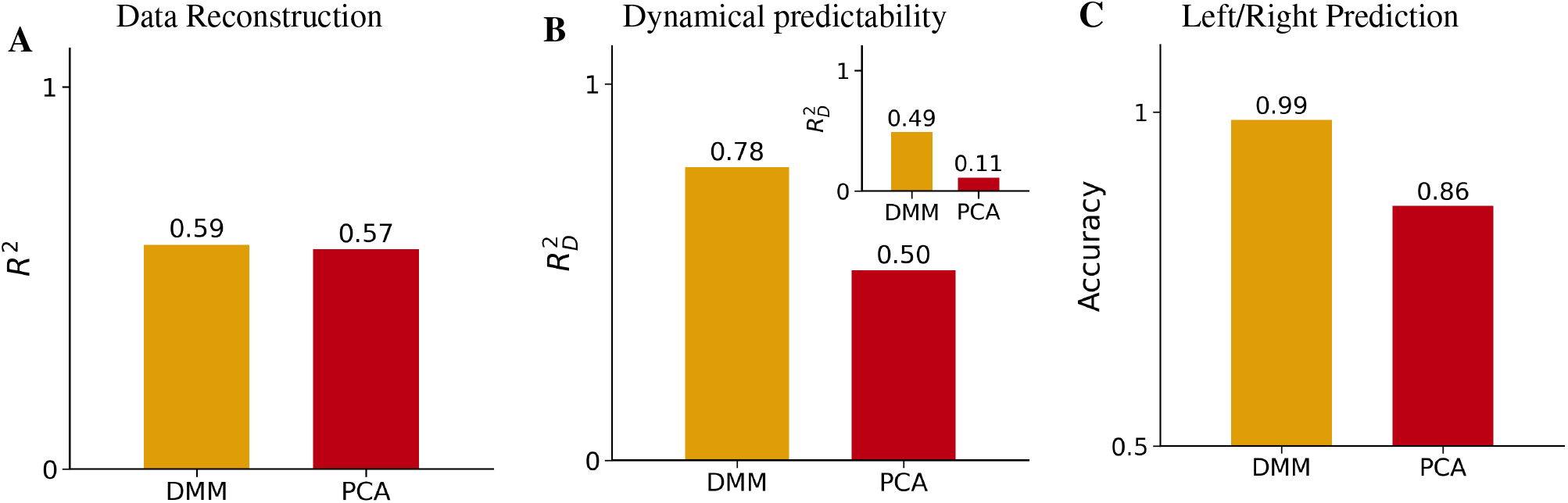
The DMM captures genuine dynamics, PCA does not. A model that learns dynamical rules (DMM) captures the temporal evolution of PMd activity, unlike a model that only compresses spatial patterns (PCA). Comparison between 3D PCA and 3D DMM. (A) Data Reconstruction (*R*^2^) shows they perform very similar in capturing the static structure of the neural trajectories. (B) Dynamical predictability (how well the latent state at one moment predicts the next). We trained a Non-Markov model to predict *z*_*t*+1_ from *z*_1:*t*_. The dynamical 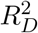 is significantly higher for the DMM, showing it has learned a more coherent dynamical structure. Inset: The Markovian predictor (*z*_*t*_ → *z*_*t*+1_) also shows higher predictability for DMM (0.49) compared to PCA (0.11). (C) Left/Right prediction accuracy. We trained a classifier to predict the trial direction given the entire latent trajectory *z*_1:*T*_ . The accuracy is almost perfect for the DMM, while for PCA results are far from optimal.

In practice, reaction time regression and movement detection are comparatively easy decoding tasks, and both DMM and PCA representations achieve near-ceiling performance on them. A more stringent test is whether the latent space encodes the reach direction, which is instructed by the peripheral target and must be represented in PMd activity ahead of movement onset to guide execution. We therefore used direction decoding accuracy as a more sensitive measure of task-relevant information. Under this stricter criterion, the DMM achieved near-perfect discrimination between leftward and rightward motor plans (Figure 2C, accuracy = 0.99), substantially outperforming PCA (accuracy = 0.86), which proved insufficient for reliable motor plan classification.

Having established our model’s superiority at capturing dynamics, we next sought to identify the true “stiff” dimensionality (Machta et al., 2013) of this dynamical engine. To this end, we trained DMM models of increasing latent dimensionality (*D* = 2, 3, 4), matched for reconstruction accuracy, and evaluated them on the previous metrics (Fig. 3).

**Figure 3.**
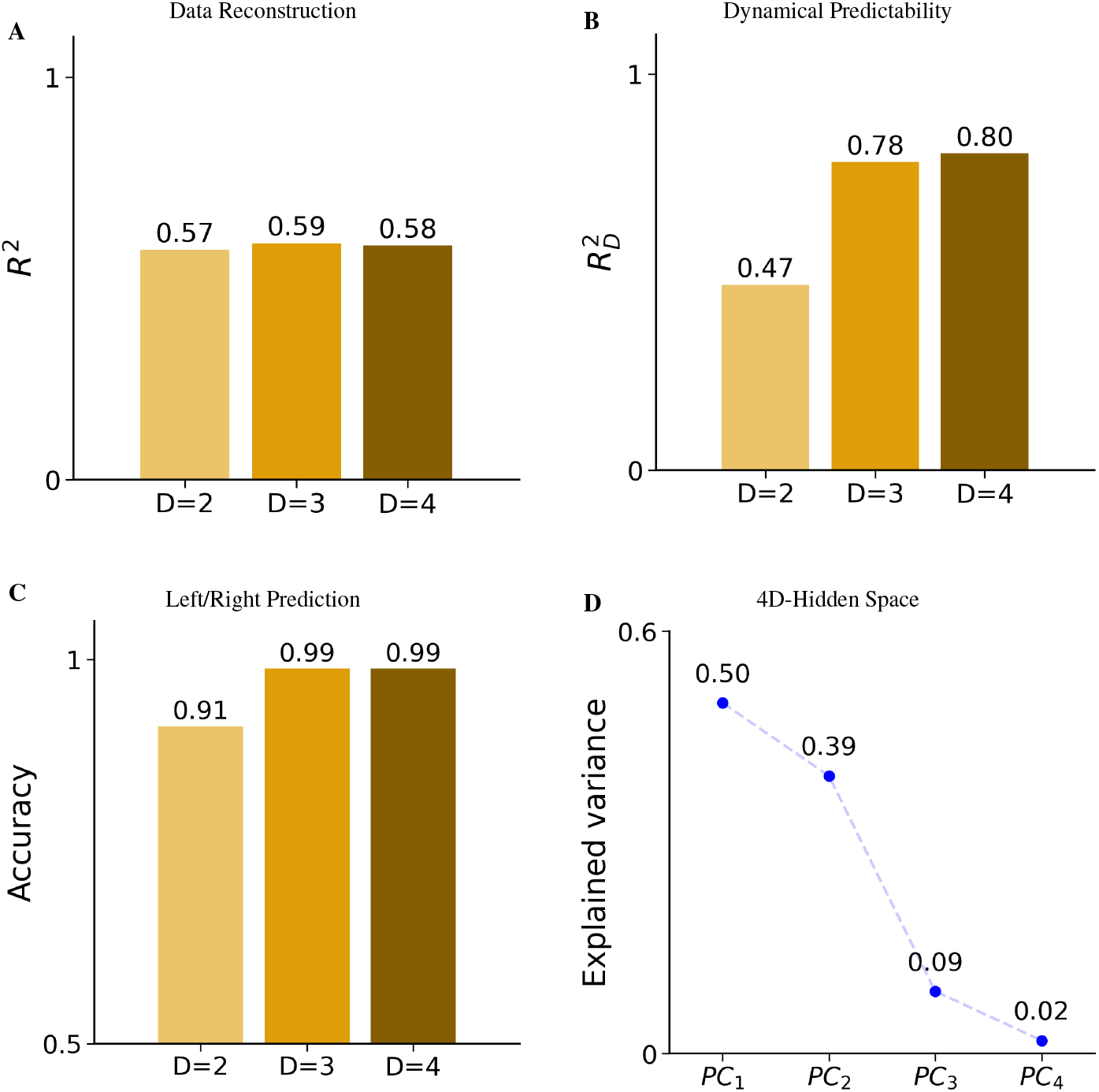
Identifying Three Dimensions as the Functional Tipping Point. A “dashboard” of metrics reveals D=3 as the minimal dimension for functional completeness. Models with latent dimensionality *D* = 2, 3, 4 were trained with matched reconstruction accuracy to isolate differences in the encoding of dynamical information. (A) Data reconstruction (*R*^2^) shows that all three models capture the static structure of the neural trajectories equally well, indicating comparable representational capacity. (B) Explained dynamical variance 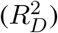 measures the ability of the latent trajectories to predict their own temporal evolution. Dynamical predictability increases sharply from *D* = 2 to *D* = 3, saturating thereafter, indicating that two latent dimensions are insufficient to capture the temporal evolution of the system. (C) Left/right prediction accuracy from the latent trajectories. While *D* = 2 provides moderate accuracy, three dimensions are required to achieve near-perfect decoding, with no improvement for *D* = 4. (D) Principal Component Analysis (PCA) of the *D* = 4 latent trajectories shows that the first three components account for nearly all variance, whereas the fourth dimension contributes negligibly, confirming that it encodes little dynamically relevant information.

This analysis revealed a critical finding. As shown in Fig. 3B, the model’s dynamical predictability 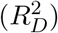 exhibits a dramatic jump from *D* = 2 (0.47) to *D* = 3 (0.78), indicating that 2 dimensions are insufficient to capture the full temporal evolution of the system. Furthermore, while a 2D model achieved reasonable decoding accuracy (Fig. 3C, 0.91), it remained suboptimal for reliably separating “Left” vs “Right” motor plans compared to the near-perfect performance of the 3D model (0.99). Crucially, increasing the dimensionality to *D* = 4 yielded negligible gains in performance. Furthermore, PCA performed on the 4D latent states confirmed that the 4th dimension accounted for very low variance (Fig 3D), indicating it contributed little to dynamical prediction. Importantly, reducing the KL regularization strength *β* by a factor of 10 (S2), did not lead to the activation of this fourth dimension, indicating that its suppression is driven by the data rather than by optimization constraints (over-pruning). Therefore, we identify *D* = 3 as the minimal “tipping point” for functional completeness.

In order to confirm this intuition, and exclude the possibility that it is just over-pruning, in the next section we further tested the generative capabilities of the DMM.

### 2.3 The 3D engine emergently predicts the full reaction-time distribution

The 3D engine, trained only on neural activity, emergently reproduced the animal’s full distribution of reaction times (Fig. 4A). To generate this prediction, we rolled out the stochastic transition model forward in time from the Go cue of all test trials using, for each trial, the same left/right context signal *c*_*t*_ observed in that trial, so that the simulated trajectory follows the same instructed condition as the real one. Afterwards, we passed the resulting latent trajectories through the RT detector (Sec. 4.4) to obtain predicted reaction times, then pooled across both reach directions, consistent with the aggregate (direction-agnostic) behavioural RT distribution against which the model is compared.

**Figure 4.**
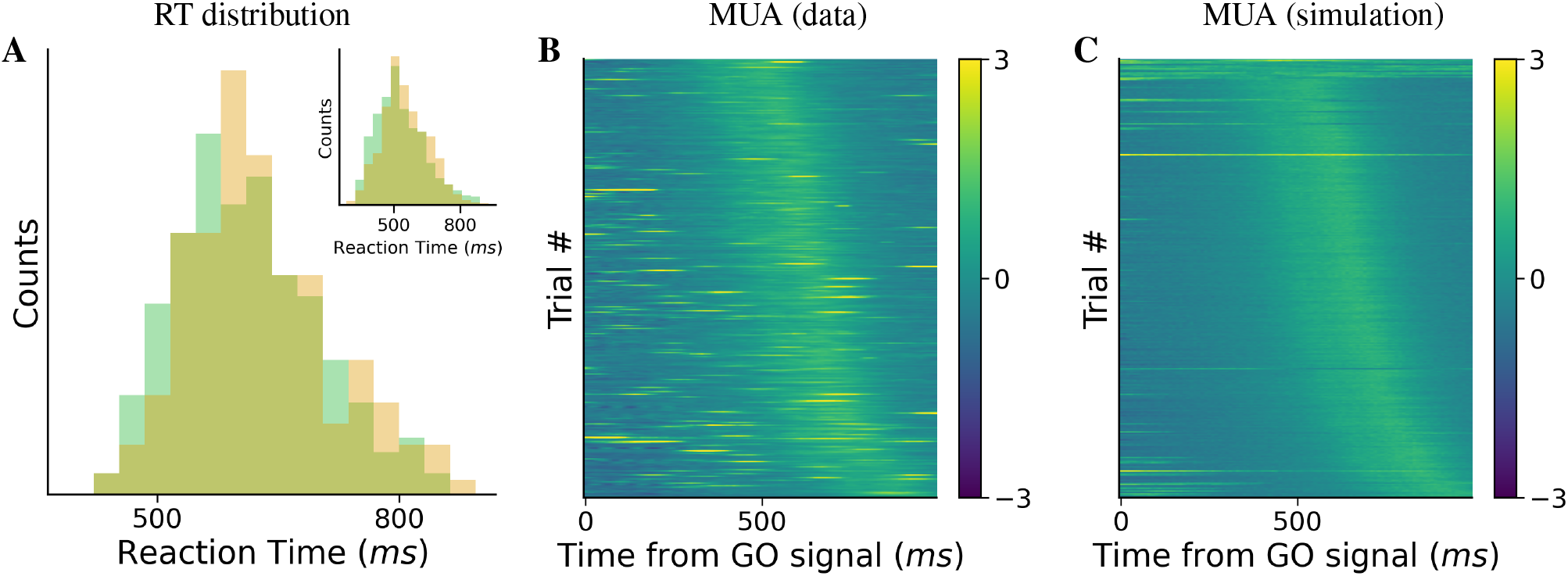
Zero-Shot Prediction of behaviour from Neural Dynamics. **(A)** The 3D model emergently generates the behavioural reaction time distribution. The histogram of the animal’s observed behavioural RTs (green) is overlaid with the histogram of simulated RTs (amber) generated by the 3D model. The simulated RTs were produced by: (1) generating trajectories starting from the GO of the inferred ones and (2) passing these trajectories through the RT detector that outputs the simulated RTs. The model was trained only on neural data, making this distribution a true, zero-shot prediction of the behavioural interface. The simulated distribution closely reproduces the overall shape of the observed RT distribution: two-sample Kolmogorov–Smirnov tests yielded *D* = 0.1104, *p* = 3.75 × 10^*−*2^ for Piero (*n* = 326) and *D* = 0.1111, *p* = 5.0 × 10^*−*4^ for Cornelio (*n* = 666). **(B)** Mean multi-unit activity (averaged across channels) for all test trials, sorted by reaction time. **(C)** The corresponding activity reconstructed from the DMM-generated latent trajectories. The simulated activity reproduces the characteristic features of the data across the RT-sorted trial axis.

Before examining the behavioural prediction, we verified that the generated trajectories reproduce the structure of the underlying neural activity. Fig. 4B shows the mean multi-unit activity (averaged across channels) for all test trials, sorted by reaction time; Fig. 4C shows the corresponding activity reconstructed from the DMM-generated latent trajectories, sorted in the same way. The two heatmaps are visually matched: both exhibit the characteristic ramping following the Go cue, with the same spatiotemporal pattern across the RT-sorted trial axis. This confirms that the generative engine faithfully recapitulates the temporal structure of the recorded neural population, not merely in aggregate but across the full range of behavioural variability.

The predicted RT distribution for Monkey P (Fig. 4A) closely matches the animal’s actual behavioural RT distribution. Importantly, this generative capacity replicates across subjects; for Monkey C (Fig. 4A, inset), the model successfully recovers the correct behavioral regime, accurately capturing both the distribution’s characteristic range and its heavy tail. Since the generative dynamics were learned only from the neural data — and the RT detector was trained on inferred latent trajectories, not on generated ones — the predicted RT distributions are not fits to behavioural data but genuine emergent predictions: the engine was never optimized, directly or indirectly, to reproduce any behavioural quantity. This cross-subject replication highlights a critical finding: across different subjects, the intrinsic, stochastic dynamics of a 3D manifold in PMd are, by themselves, sufficient to generate the full spectrum of behavioural variability.

This predictive power extends to the single-trial level, consistent with evidence that latent dynamics inferred via nonlinear sequential models track behavioural output on a moment-by-moment basis (Pandarinath et al., 2018). To quantify trial-specific correspondence between latent dynamics and behaviour, we computed, across a broad range of simulation start times, the correlation across trials between the true reaction times and the reaction times obtained by rolling out each trial’s latent state from the corresponding time point, using the appropriate left/right context signal. Figure 5 shows how this correlation increases as the simulation start time approaches movement onset.

**Figure 5.**
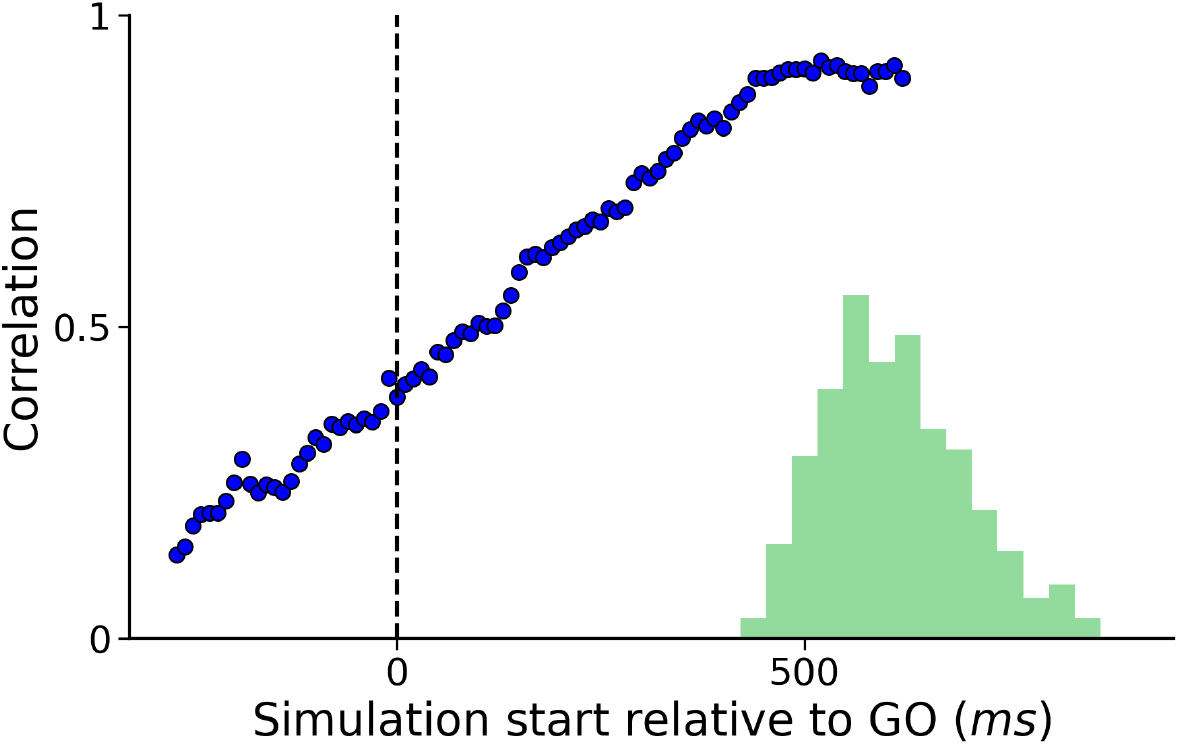
Latent Dynamics Constrain Action Timing on Single Trials. Temporal evolution of single-trial predictive power. The line graph shows the correlation (Y-axis) between the animal’s actual RT and the model’s predicted RT as a function of the simulation start time (X-axis), relative to the GO cue. The correlation increases as the simulation incorporates more neural data closer to movement onset, demonstrating that the latent dynamics increasingly constrain the specific timing of the action. The black vertical dashed line marks the Go cue, where the correlation is already substantial (approximately 0.4). For reference, the distribution of true reaction times, the same of Fig. 4A, is shown in the lower right of the figure, providing a visual indication of the temporal proximity between each simulation start point and the behavioural event being predicted.

Notably, even when rollouts are initialized at the Go cue (vertical dashed line), the model achieves a non-trivial correlation with the true reaction times (*r* ≈ 0.4). This indicates that the latent state inferred at the Go cue already contains meaningful information about upcoming behavioural variability, despite the absence of overt movement. At the same time, the moderate magnitude of this correlation reflects intrinsic stochasticity in the latent dynamics, which progressively differentiates trials over time and limits the predictability of reaction time from early trial states alone. As the rollout initialization moves closer to the reaction time, an increasing portion of the trial-specific dynamical evolution has already unfolded, reducing uncertainty and improving prediction accuracy.

Although prediction accuracy improves as rollouts are initialized later in the trial, the correlation does not approach unity, even at late time points, consistent with residual noise in both the latent dynamics and the behavioural readout.

Collectively, these findings demonstrate that the DMM identifies stochastic, behaviourally informative latent dynamics, while confirming that the pruned dimension is non-informative, establishing three as the minimal intrinsic dimensionality of the system.

### 2.4 The neural state at the Stop signal predicts inhibitory outcome

Up to this point, our analysis focused exclusively on no-stop trials. We now extend the framework to stop trials, distinguishing between successful stops, in which the monkey inhibits movement following the Stop signal, and unsuccessful stops, in which the Stop signal is unable to prevent movement initiation.

We first examined how these two trial types are represented in the inferred latent space. For visualization purposes, all latent-space plots show the projection onto the first two latent components. Fig. 6A shows representative latent trajectories from a successful stop trial (red) and an unsuccessful stop trial (green). In both cases, trajectories initially depart from the pre-Go baseline following the Go cue. After the Stop signal, however, their dynamics diverge sharply. Successful stop trials exhibit a rapid reversal, returning toward the pre-Go region of latent space. A notable geometric distinction is visible: whereas no-stop trajectories trace broad, extended orbits through the latent space, successful stop trajectories are deflected toward more internal, shorter-radius orbits—a compression consistent with the arrest and reversal of the motor plan before full execution. In contrast, unsuccessful stop trials are minimally perturbed by the Stop signal and continue along a curved trajectory toward movement-related regions, only returning toward baseline after movement onset.

**Figure 6.**
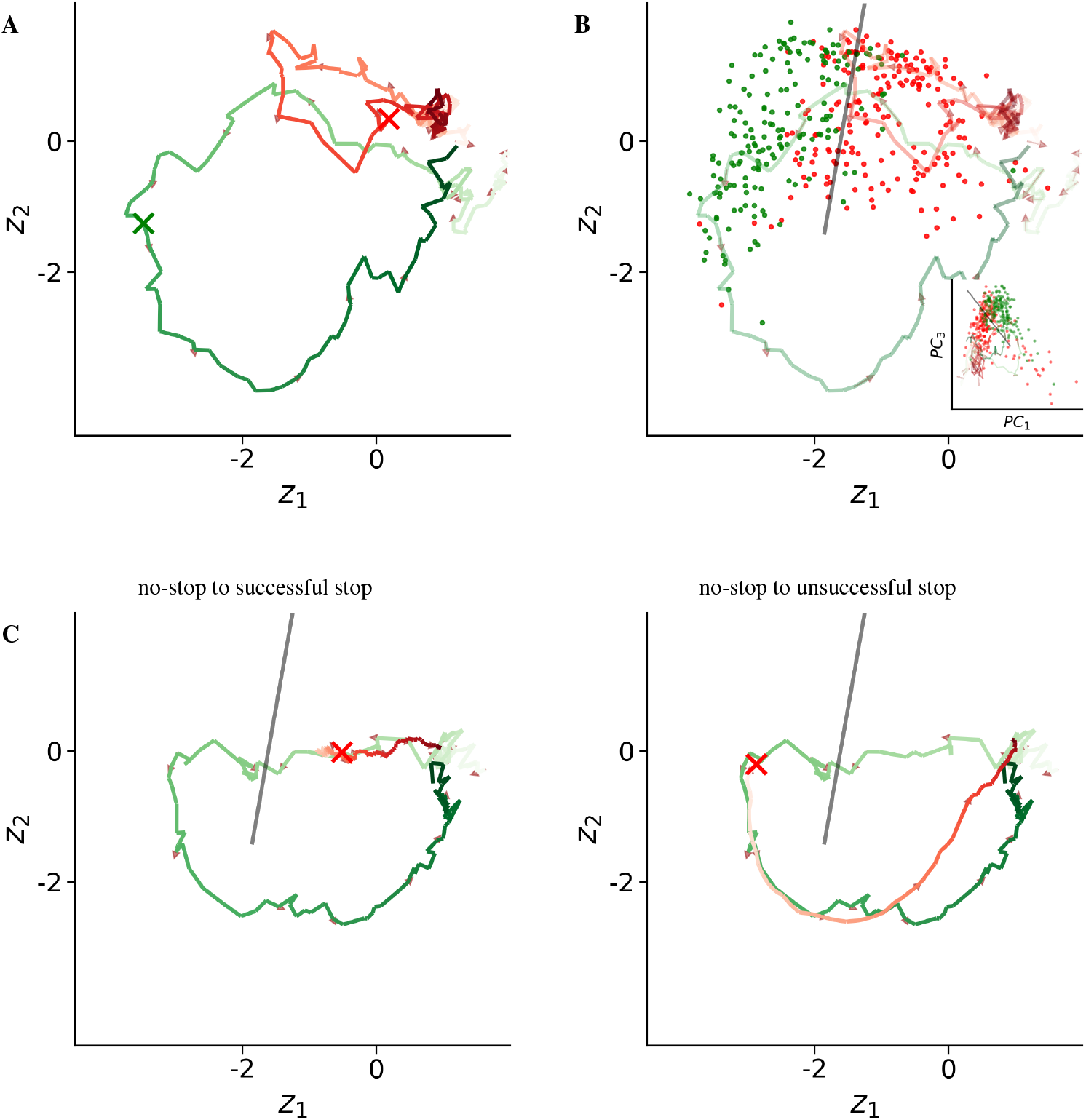
State-dependent stopping dynamics in latent space. **(A)** Representative latent trajectories for a successful Stop trial (red) and an unsuccessful Stop trial (green), projected onto the first two latent dimensions (color gradient indicates temporal progression). Following the Stop signal (cross), successful Stop trajectories reverse toward the pre-Go region of latent space, whereas unsuccessful Stop trajectories continue toward movement-related regions before returning to pre-GO after movement onset. **(B)** Neural State at Stop Signal Predicts Inhibition Success. Latent states at Stop-signal time for all trials (successful Stops in red, unsuccessful Stops in green). The black line indicates the linear discriminant boundary, achieving approximately 86% discrimination accuracy, compared to a lower 80% accuracy in the PC space (inset), on the training set. Partial overlap reflects intrinsic stochastic variability. **(C)** Simulations (red) generated from the same inferred no-stop trajectory (green) under different virtual Stop-Signal Delays (SSD); the simulated trajectory coincides with the no-stop trajectory up to the injection point, after which the two diverge. Left: early Stop injection (short SSD) causes the generated trajectory to deviate toward baseline, producing a successful Stop. Right: late Stop injection (long SSD) allows the trajectory to continue toward movement-related regions, resulting in an unsuccessful Stop.

To test whether this divergence reflects a systematic organization of stopping outcomes, we plotted the latent states at the time of the Stop signal for all training trials (Figure 6B); we used the training set specifically because it offers a substantially larger number of trials than the test set alone, where the small number of Stop trials would make an accuracy estimate statistically unreliable, making the separation structure easier to characterize robustly. This revealed a partial but systematic separation between successful and unsuccessful stops. A linear discriminant boundary trained on these states achieves an accuracy of approximately 86%, indicating that stopping outcome is strongly, though not perfectly, encoded at the time of the Stop signal. Given the low capacity of a linear classifier relative to the number and dimensionality of the training points, overfitting is unlikely, so this accuracy is expected to reflect the genuine separability of the two outcome classes rather than memorization of the training set. When the same classifier is trained on the first three principal components obtained by PCA, its accuracy drops to 80%. The imperfect separability reflects intrinsic trial-to-trial variability and stochastic latent dynamics following the Stop signal.

We next asked whether the learned dynamics support mechanistic manipulation of stopping behaviour. Starting from inferred no-stop trials, we injected a virtual Stop signal at different time points and generated the resulting trajectories under the Stop context. Fig. 6C illustrates two such in silico simulations using the same underlying Go trial. When the Stop signal is applied early (short SSD; left), the generated trajectory diverges from the original Go dynamics and reverses toward the baseline region of latent space, leading the movement outcome detector to classify the trial as a successful stop. When the Stop signal is applied later (long SSD; right), the generated trajectory closely follows the original Go dynamics toward movement-related regions and is classified as an unsuccessful stop.

Crucially, both outcomes arise from the same initial trial and differ only in the latent state at which the Stop context is engaged. Together, these results show that the DMM captures a continuous, state-dependent stopping process, in which inhibitory success emerges from the interaction between ongoing latent dynamics and the timing of the Stop signal, rather than from a race between mutually independent processes.

### 2.5 In silico experiments reveals systematic violations of the IRM’s independence axioms

With the 3D engine validated as a generative model of the Go process, we can now test if the neural manifold permit the Go and Stop processes to operate independently, as assumed by the IRM. The latter rests on two formally distinct assumptions (Logan and Cowan, 1984; Verbruggen et al., 2019): context independence, which requires that the Go reaction-time distribution be identical on stop and no-stop trials, and stochastic independence, which requires that the finishing times of the Go and Stop processes, respectively the RT and the SSRT, are statistically independent. We test each in turn, beginning with context independence.

#### 2.5.1 Stop-failure reaction times violate context independence

We asked whether the stop process, when it fails to prevent a response, leaves the Go reaction time unperturbed—the defining requirement of context independence. To test this, we used the generative engine to reconstruct a counterfactual: the reaction time each Go trial would have produced had a stop signal been presented and failed. For each Go trial, we injected a virtual stop signal at a given SSD into the inferred latent trajectory, simulated the subsequent dynamics under the Stop context, and—for simulations in which the stop failed—estimated the resulting reaction time (see Sec. 4.7 for the full procedure). The difference ΔRT = RT_stop failure_ − RT_Go_, where RT_Go_ is the trial’s own behavioural Go reaction time, measures how much the stop process distorts the Go finishing time. Under context independence, ΔRT should be zero at every SSD. This per-trial construction is also an advantage over behavioural approaches such as (Bissett et al., 2021b), which must contrast stop-failure RTs from one set of trials against Go RTs from a disjoint set. By pairing each simulated stop-failure RT with the *same* trial’s own Go RT, our counterfactual removes the across-trial pairing variance that otherwise inflates estimates of the stop-induced RT distortion. Anchoring to the trial’s real Go RT is well-justified here: per-trial simulated and true RTs are already strongly correlated early in the trajectory (Fig. 5), so the real RT is a faithful baseline for the matched stop-failure simulation.

Instead, the model predicts a rich, SSD-dependent structure (Fig. 7). For Monkey P (main panel), ΔRT is strongly positive at short SSDs—stop-failure responses are slower than the corresponding Go responses—and crosses zero to become slightly negative at longer SSDs, where stop-failure responses become marginally faster. This non-monotonic pattern closely mirrors the human behavioural data reported by Bissett et al. (2021b), who documented the same qualitative signature across a large sample of participants performing the stop-signal task. That our model, trained exclusively on macaque premotor neural activity, spontaneously reproduces this behavioural signature from its intrinsic dynamics represents an independent, mechanistic confirmation of the Bissett et al. (2021b) finding and demonstrates that the violation originates in the neural implementation itself.

**Figure 7.**
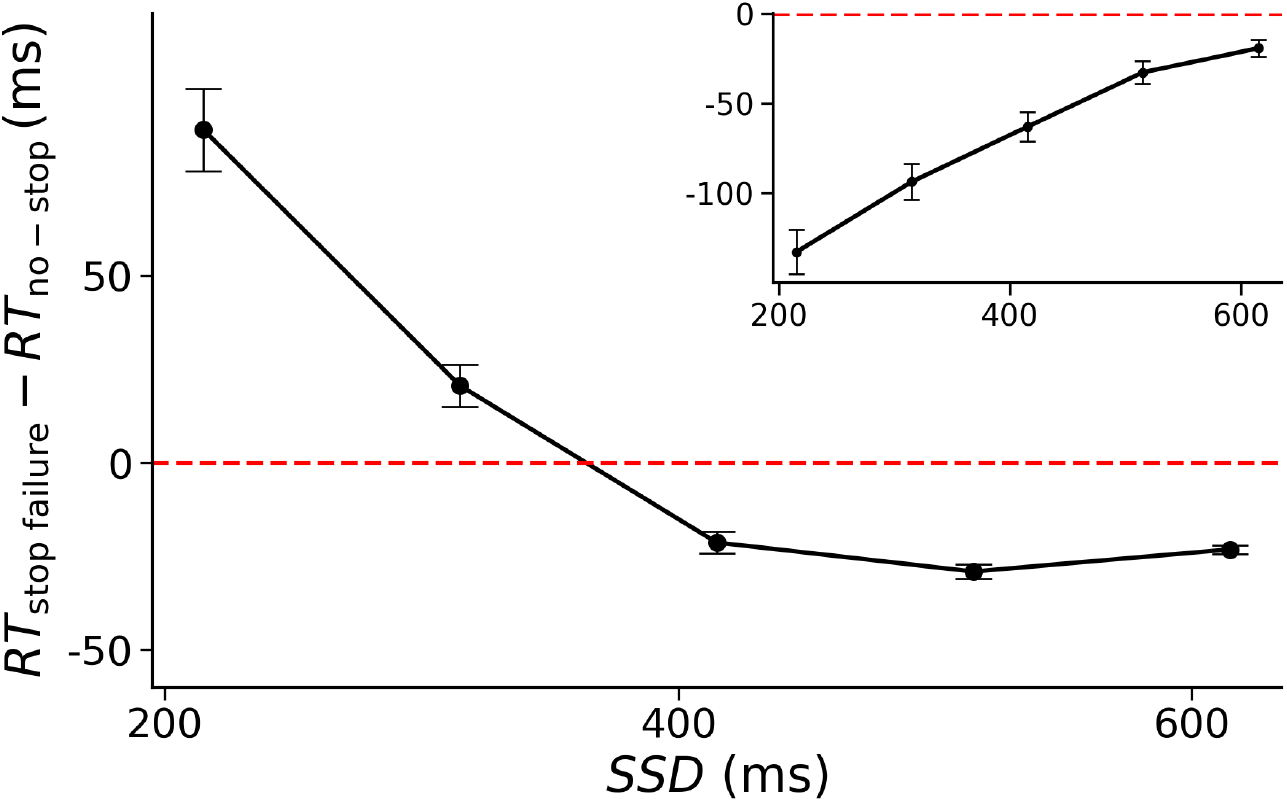
The underlying dynamics reveal a violation of context independence. Mean ΔRT = RT_stop failure_ − RT_Go_ as a function of stop-signal delay (SSD). For each Go trial, a virtual stop signal was injected and the resulting stop-failure RT was estimated via simulation (see Sec.4.7). Error bars: SEM. The red dashed line marks ΔRT = 0, the prediction of context independence. Main panel (Monkey P): ΔRT is positive at short SSDs and slightly negative at long SSDs, closely mirroring the pattern reported by Bissett et al. (2021b). Inset (Monkey C): ΔRT is negative at all SSDs, with larger deviations at shorter delays. Both profiles violate context independence.

For Monkey C (Fig. 7, inset), ΔRT is negative at all SSDs, with increasingly negative values at shorter SSDs. Although this pattern differs from Monkey P’s, it constitutes an equally clear violation of context independence: the stop process accelerates stop-failure responses, and does so more strongly the earlier it intervenes. Both patterns point to the same fundamental conclusion: the stop process does not leave the Go process undisturbed. The shared neural manifold compels interaction.

#### 2.5.2 Per-trial SSRT analysis violates stochastic independence

Having shown that context independence does not hold at the level of the underlying neural dynamics, we next examined the second pillar of the IRM, *stochastic independence*: the requirement that, on any single trial, the finishing times of the Go and Stop processes be uncorrelated. Our in silico framework lets us “run the race” trial by trial (Fig. 6): for a given Go trial’s neural trajectory, we inject a Stop context at a chosen Stop-Signal Delay (SSD) and pass the perturbed trajectory through the movement detector (Sec. 4.4), which returns the probability of movement, *p*(move). By locating the critical SSD at which this classification flips, we obtain a mechanistic, per-trial SSRT. We use a per-trial estimator here, distinct from the population-level integration method used for the cross-species comparison of Fig. S6, for a specific reason: testing stochastic independence requires correlating each trial’s RT with its own SSRT, and the integration method is structurally aggregate—it returns a single SSRT for the whole distribution and cannot, by construction, yield a per-trial value. We then ask whether, across trials, the Go process finishing time (indexed by RT) predicts the Stop process finishing time (indexed by SSRT). It does: the simulation reveals a strong *positive* correlation between a trial’s RT and its SSRT (Fig. 8), in direct contradiction of stochastic independence, and the effect is replicated in the second animal (Monkey C, Fig. 8, inset).

**Figure 8.**
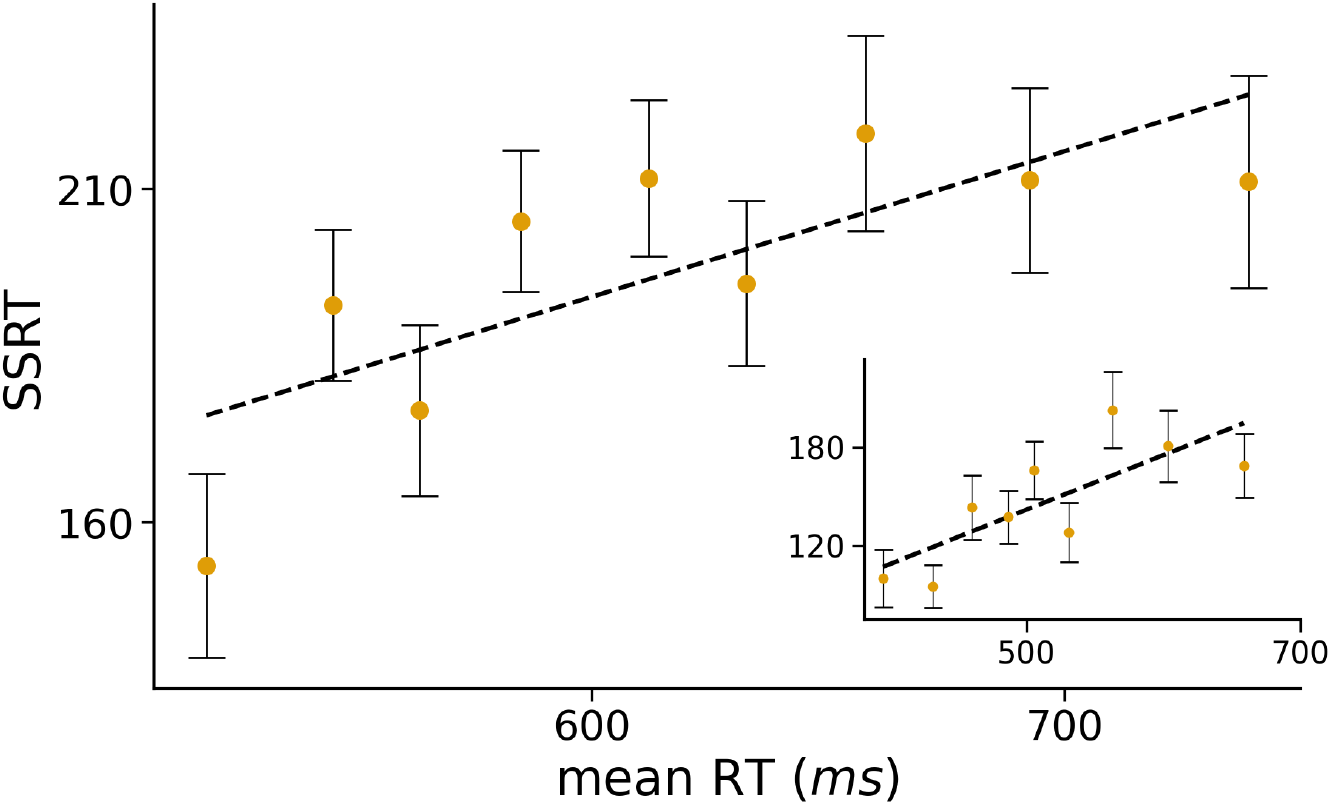
The underlying dynamics reveal a violation of stochastic independence. Mean SSRT as a function of mean RT, computed by grouping trials into nine equally sized RT bins (error bars: SEM). The black dashed line shows a linear regression fit. The simulation reveals a strong positive correlation between the mean RT of a trial type and its estimated SSRT, exposing a dependence between the Go and Stop processes that the Race Model’s independence assumption does not capture.

Because our per-trial SSRT is defined very differently from the familiar behavioural estimate, its logic warrants a brief clarification—particularly to avoid a natural misreading. The standard integration method (Logan and Cowan, 1984; Verbruggen et al., 2013) derives a single, population-level SSRT from the aggregate Go RT distribution: it rank-orders no-stop Go RTs, takes the *n*th RT (with *n* set by the observed probability of responding on stop trials), and subtracts the SSD—a procedure rooted in the Independent Race Model, which itself assumes both forms of independence. Our procedure never consults the RT distribution.

We simulated each trial forward, under the Stop context, from a dense range of virtual stop times, generating multiple stochastic rollouts at each and estimating *p*_Move_ as the fraction classified as movements by the movement classifier (see Sec. 4.4). The per-trial critical SSD, *SSD*^∗^, is defined as the latest virtual stop time at which *p*_Move_ remains below 0.5, and the per-trial SSRT is computed as *RT* − *SSD*^∗^, where RT is the trial’s observed reaction time. This 0.5 criterion follows from the definition of SSRT as the time the stop process takes to complete inhibition: if the stop signal arrives exactly one SSRT before the RT, Go and Stop are expected to finish at nearly the same moment, so a stop delivered at that instant should be inhibited in about half of the simulated futures, i.e *p*_Move_ ≈ 0.5. This is distinct from the 50% population stopping rate of the experimental staircase: the staircase’s 50% is a fraction of trials, ours is the fraction of simulated futures from a single inferred state, thus an averaged proxy rather than a faithful per-trial estimate. Because of this, trials are subsequently pooled into groups and analysed via group-level mean RT and SSRT (see Sec. 4.4) rather than individually. Also, because our SSRT is read out from each trial’s own neural dynamics rather than from the aggregate Go RT distribution, RT and SSRT cannot be circularly bound to a shared distributional assumption, making the observed correlation a genuine test of independence rather than an artifact of estimation.

A second objection concerns the model class itself: could the RT–SSRT dependence simply be an artifact of a nonlinear architecture in which all latent dimensions interact? Nothing in the DMM forces such interaction. A 2-dimensional DMM could trivially implement the IRM—one latent dimension for Go, one for Stop, with the learned dynamics holding them independent. That the data-driven model instead settles on interactive dynamics over a shared 3D manifold is a property of the neural data, not of the model class. Nonlinearity, moreover, is a capacity rather than a requirement: had the true dynamics been linear, the transition function could have approximated them. The model’s flexibility makes the observed coupling *more* compelling, not less, because the model was free to learn independence and did not.

This dependence is therefore not a parameter choice but an emergent property of the ‘stiff’ neural dynamics, conserved across both animals. We stress that this is not simply a claim that our model fits the data better than the IRM. It is a mechanistic claim: the model—a validated proxy for the neural hardware—shows that the strict independence the Race Model requires at the algorithmic level is incompatible with the sufficient conditions for generating the behaviour (the 3D neural manifold). At the level of the underlying implementation, the biological substrate simply lacks the degrees of freedom to support two fully independent processes—even as the IRM continues to offer an accurate and parsimonious description of behaviour itself.

The positive *sign* of this dependence deserves emphasis. Colonius and Diederich (2018) showed that a race model with perfect negative dependence preserves mean SSRT estimates, offering a theoretical resolution of the independence paradox. Our manifold-constrained dynamics instead produce positive dependence—faster Go implies faster Stop—consistent with single-trial EMG evidence that go- and stop-process latencies covary positively within individual trials (Raud et al., 2022; Jana et al., 2020). Colonius et al. (2024) further showed that behavioural data alone cannot distinguish context from stochastic independence violations; our neural-level approach bypasses this identifiability problem entirely.

Averaged across all trials, our per-trial procedure recovers a population SSRT (≈ 190 ms) almost identical to the integration-method estimate (≈ 183 ms), highlighting an important subtlety: the IRM’s *aggregate* behavioural predictions remain approximately correct even though independence does not hold at the level of the underlying dynamics. This is precisely why the IRM has remained such a useful and durable behavioural model: the aggregate statistics it predicts are largely insensitive to the underlying dependence structure, as Colonius and Diederich (2018) anticipated. What our results add is the mechanism beneath that success: independence is not merely unsupported by the data but structurally precluded by the geometry of the manifold from which the data emerge, a layer of detail the IRM, operating on behaviour alone, could not have accessed.

These results can be read alongside previous population-level analyses of stopping. Pani et al. (2022) showed that in PMd the neural trajectories of successful-stop and latency-matched no-stop trials diverge approximately 90 ms after the Stop signal—well before the behavioural SSRT of ≈ 190 ms—an interval interpretable as the time for the Stop signal to become effective at the population level. SSRT thus comprises two stages: an initial interval separating the Stop signal from the divergence of successful-stop and no-stop trajectories, and a subsequent interval in which the trajectory progresses from leaving the output-null subspace to the commitment point beyond which movement can no longer be prevented. Consistent with this, latent states at the time of the Stop signal separate successful from unsuccessful stops with an accuracy of approximately 86%, limited by the stochastic dynamics governing the first interval.

Within this framework, the dependence of SSRT on RT arises naturally. Trials with longer RTs reflect slower or noisier latent dynamics, which on average take longer to progress from the Stop signal to the commitment threshold; faster trials reach it sooner, yielding shorter SSRTs. While variability in the latency of stop-signal effectiveness may contribute at the single-trial level, the dependence of SSRT on RT emerges robustly at the population level.

### 2.6 Targeted neural perturbations systematically shift reaction times in silico

Finally, we used the model as a “virtual brain” to design and test targeted perturbation protocols. Because the perturbations are performed on the multi-unit activity, they constitute experimentally testable predictions, echoing in vivo work in which subthreshold intracortical microstimulation of PMd during the delay period was shown to causally delay movement onset without disrupting the movement itself (Churchland and Shenoy, 2007). Here, we asked whether an analogous causal link could be recovered directly from the model. Indeed, targeted stimulation of MUA produced robust, sign-specific shifts in predicted reaction times, consistently accelerating or delaying movement onset depending on the polarity of the applied perturbation (Fig. 9).

**Figure 9.**
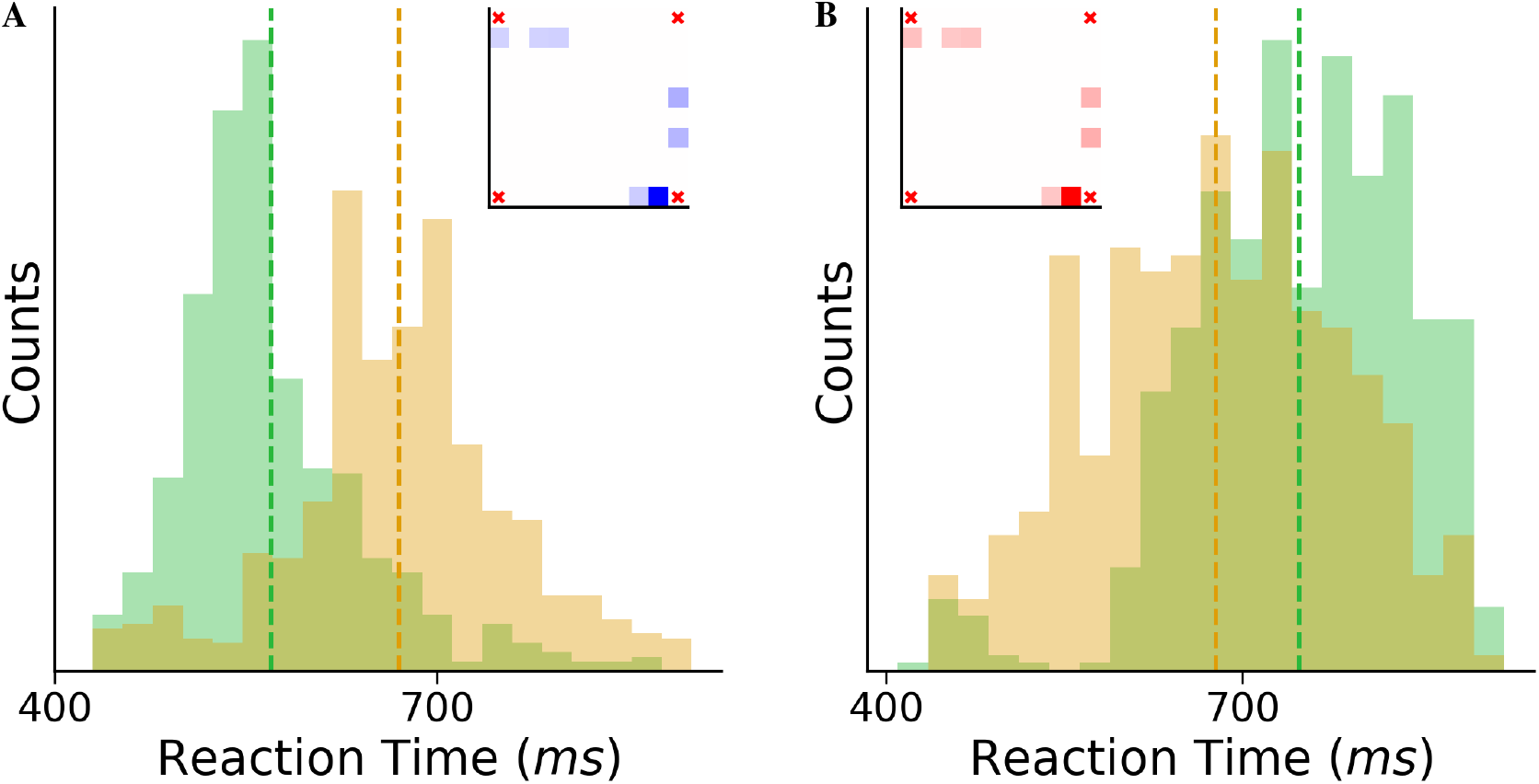
In silico intervention: Causal experiments in a “virtual brain”. Virtual stimulation applied directly on the normalized multi-unit activity induces systematic and polarity-dependent shifts in predicted reaction times. (A) Lengthening fast trials. The RT distribution of the 10% of trials with the shortest RTs, before (green) and after (amber) stimulation. The shift to the right indicates a systematic slowing of the movement plan. (B) Shortening slow trials. The RT distribution of the 10% of trials with the longest RTs, before (green) and after (amber) stimulation. The shift to the left indicates a systematic acceleration of the movement plan. Vertical dashed lines indicate the corresponding distribution means. Stimulation produces substantial and asymmetric RT shifts (approximately +100 ms and − 80 ms), demonstrating a mechanistic influence of the perturbed neural activity on behaviour. *Inset:* Last mean MUA perturbation Δ*x*_*GO*+100 ms_, averaged across trials, shown as a spatial map over the electrode array (blue refers to negative values, while red to positive). For visualization, electrodes with stimulus amplitude below 0.05 standard deviations (SD) in standardized MUA units, were thresholded to zero, revealing a sparse pattern involving only 6–7 electrodes. Perturbation amplitudes across electrodes remain small, with the largest reaching ∼ 0.3 SD, still below the intrinsic per-step variability of the neural activity, indicating that the applied stimulation is biologically plausible and, in principle, reproducible in real experiments. Insets reveal that the stimulation patterns for RT lengthening and RT shortening are approximately mirror images — opposite in sign across the same sparse set of electrodes — confirming that the perturbation acts along a single direction in neural space.

Perturbations aligned with the RT-increasing latent direction (effectively navigating the intrinsic manifold, see Jazayeri and Afraz (2017)) resulted in a pronounced rightward shift of the RT distribution, with a mean delay of approximately 100 ms, while stimulation in the opposite direction yielded a leftward shift of approximately 80 ms. Crucially, control experiments with infinitesimal perturbations produced overlapping RT distributions between stimulated and unstimulated simulations, ruling out the possibility that the observed effects arise from a generic regression-to-the-mean behaviour of the DMM when simulating short- or long-RT trials. Instead, the shifts reflect a genuine mechanistic consequence of the applied stimulus.

The mean stimulation pattern at the final time step (*GO* + 100*ms*), shown as an inset in Fig. 9, reveals that only a small subset of electrodes (6–7 channels) received substantial modulation. For visualization, electrodes with stimulus amplitude below 0.05 standard deviation (SD) in standardized MUA units, were thresholded to zero, as such values are negligible compared to the intrinsic per-step activity variability (SD ≈ 0.4; Supplementary Fig. S4C). Moreover, the maximum stimulus amplitude per electrode (∼ 0.3 SD) remains below this variability scale, indicating that the applied perturbations are both realistic and experimentally feasible.

Together, these results demonstrate that the DMM generates precise, experimentally testable predictions for how targeted neural stimulation should shift reaction times — predictions that can be directly verified using intracortical microstimulation in future experiments.

## 3 Discussion

Our central finding is that the motor dynamics of the macaque premotor cortex can be captured by a compact, 3-dimensional Markov process that, by itself, generates the full spectrum of behavioural reaction times. This engine reveals that the Go and Stop processes are not independent but are coupled by the geometry of a shared neural manifold — a mechanistic layer of structure that the Independent Race Model, operating on behaviour alone, could not access.

Crucially, this rejection does not invalidate the IRM’s aggregate behavioural predictions, which remain approximately correct; rather, it replaces an abstract computational assumption with a physical mechanism, providing the generative Algorithm (Marr and Poggio, 1976) that connects the neural implementation to behavioural dynamics.

Our findings provide this missing link. The core of this argument rests on a new standard for model selection: rather than maximizing explained variance, we required the model to be functionally complete — capable of hosting all computations the task demands, including the separation of competing motor plans. By this criterion, three dimensions constitute the minimal “stiff” manifold (Gutenkunst et al., 2007; Machta et al., 2013), while two are insufficient and four are redundant.

A critical but easily overlooked aspect of this finding concerns the Markov property of the dynamics. Low dimensionality alone does not guarantee a compact dynamical system: a 3-dimensional process with long temporal memory would be low-dimensional only in appearance, since its effective degrees of freedom would scale with the memory depth. The DMM’s strong one-step predictability 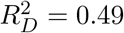 for the Markov predictor, compared to only 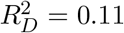 for PCA) confirms that the 3D dynamics we identify are approximately Markov. The current state of the system largely determines its next step. This means that three numbers, measured at a single moment, carry most of the information needed to predict the system’s immediate future—a far stronger claim than saying that three principal components capture most of the variance.

This 3D generative engine allowed us to bridge the gap from neurons to behaviour in a way descriptive models cannot. We showed that the model, trained only on neural data, emergently recreated the animal’s complete behavioural RT distribution—a true “zero-shot” prediction that validates it as a “functionally sufficient” model of the “Go” process. This achievement is more than a fitting exercise; it is a demonstration that the intrinsic, 3D stochastic dynamics of the PMd are the engine of behavioural variability.

One might naturally object that PMd is only a single node within a highly distributed network governing motor decisions and inhibitory control, a network that heavily involves the basal ganglia and other prefrontal cortices. Observing only PMd could thus be perceived as a limitation. However, our model’s ability to emergently reconstruct the full, complex distribution of behavioural reaction times provides a powerful counter-argument: the behaviorally relevant computations of this wider network are effectively funneled through the PMd. In this view, the PMd dynamics provides a low-dimensional representation of them. The complex processes of upstream regions are projected onto the PMd manifold, and our generative engine successfully captures this integrated net effect. Thus, while other areas are undoubtedly involved in the decision process, their behaviorally relevant contributions are largely inscribed within the local PMd dynamics we model.

By validating this 3D model as a generative replacement for the “Go” process, we were able to use it as an in silico “virtual brain” to test, at the mechanistic level, the independence assumptions of the IRM. This two-pronged simulation provided quantitative evidence of systematic violations of both of the IRM’s independence assumptions.

First, a simulated stop-failure analysis (Fig. 7) revealed SSD-dependent distortions of the Go reaction-time distribution consistent with a violation of context independence: for Monkey P, the ΔRT profile closely mirrors the non-monotonic pattern documented in humans by Bissett et al. (2021b), while Monkey C exhibits a distinct but equally clear violation. Then, a per-trial SSRT analysis (Fig. 8) further revealed a positive correlation between RT and SSRT, indicating a violation of stochastic independence—a dependence whose positive sign distinguishes our result from the negative-dependence resolution of (Colonius and Diederich, 2018) and aligns with the interactive race account (Boucher et al., 2007). Together, these results show that a single, interactive, low-dimensional system underlies the observed behaviour, coupling Go and Stop to a degree that a purely independent-racers description cannot capture.

Finally, we demonstrated how this generative model serves as a new platform for causal experimentation. By computationally perturbing the latent state (or the structured noise we identified), we can systematically and quantitatively predict the resulting change in behaviour (e.g., RT). This moves beyond the “which-neurons-do-I-stimulate” problem of spiking networks (Histed et al., 2009)—where physical stimulation activates sparse, distributed populations often failing to respect the system’s intrinsic dynamics (Jazayeri and Afraz, 2017)—and provides a principled, low-dimensional “control knob” for simulating interventions at the level of the computation itself.

This “top-down” synthesis returns to the paradox we opened with: how does stable function emerge from variable components? Our answer, for the case of motor control, is that the brain compresses its variable, high-dimensional activity onto a 3-dimensional stiff manifold whose Markov dynamics are sufficient to generate behaviour. The vast majority of neural degrees of freedom are sloppy — behaviourally irrelevant — and the system’s robustness arises precisely because function depends on so few dimensions. This principle — using high-level function to identify the low-level stiff manifold — is a generalizable template for systems neuroscience. Consistent with findings that intrinsic neural manifolds constrain learnable activity patterns (Sadtler et al., 2014), it provides a path to escape the “curse of dimensionality” and the limitations of purely descriptive models, creating predictive theories of how function emerges from implementation.

Our results elevate the descriptive ‘holding’ subspace and ‘escape’ dynamics reported by Pani et al. (2022) into a generative engine, identifying the stochastic vector field that drives the system (Fig.1B). While linear methods require three dimensions per movement direction (Pani et al., 2022), our nonlinear DMM identifies a single 3D ‘stiff’ core sufficient for the entire task; notably, *z*_2_ acts as a primary axis for both directional choice and inhibitory outcome. This model refines the concept of a rigid ‘maturation threshold’ (Pani et al., 2022) into a smooth, probabilistic transition: commitment to a motor plan matures continuously over time rather than through a binary switch, as reflected in the graded, state-dependent stopping outcomes of Fig. 6B.

The contrasting context-independence profiles of the two animals (Fig. 7) invite a further connection to the Pause-then-Cancel framework (Schmidt and Berke, 2017), which posits that action-stopping relies on two sequential mechanisms: a fast, global Pause that transiently suppresses motor output, followed by a slower, goal-directed Cancel that specifically terminates the planned action (Diesburg and Wessel, 2021; Tatz et al., 2021). In the original formulation, the Pause and Cancel are temporal stages—the Pause fires first, at fixed latency after the salient event, and the Cancel follows. We propose a complementary spatial reinterpretation, in which the two mechanisms correspond to functionally distinct regions of the latent force field. This preserves the sequential logic—the trajectory encounters the Pause region before the Cancel region—but adds a crucial degree of freedom: the effect of each mechanism depends on where in the latent space the trajectory is when the stop context engages. In this view, Monkey P’s non-monotonic ΔRT profile reflects a force field with a prominent Pause region that slows trajectories at short SSDs (ΔRT *>* 0); at long SSDs, the trajectory has already traversed this region under Go dynamics—without any slowing, since the stop context was not yet active—and the dominant effect is the Cancel, which deflects the trajectory toward the shorter-radius orbits characteristic of successful stops (cf. Fig.6A), producing marginally faster RTs (ΔRT *<* 0). Monkey C’s uniformly negative profile would correspond to a force field with a minimal Pause region, where the Cancel is the only operative mechanism.

Our biological conclusions converge with recent work using recurrent neural networks as “digital twins” of PMd (Alboré et al., 2025), which independently demonstrates that motor inhibition reflects state-dependent dynamics rather than autonomous racers. Our approach differs in parsimony: by identifying three dimensions as the minimal generative core, learning the manifold directly from raw multi-unit activity — rather than from temporally agnostic PCA projections, which discard the sequential structure of the data before any dynamics are learned (a cost we quantified directly, see Fig. 2, inset) — and by explicitly parameterizing the stochastic transition rules, the DMM provides a tighter, more identifiable link between neural variability and behavioural output. In particular, reaction-time distributions emerge as first-principles predictions rather than post-hoc fits to empirical residuals.

More broadly, our approach relates to a growing family of deep latent variable models for neural population data, including LFADS (Pandarinath et al., 2018) and switching linear dynamical systems. These methods have proven powerful for inferring smooth latent trajectories from noisy spiking data. However, ensuring that such flexible models yield accurate scientific interpretations rather than merely fitting data remains a significant challenge (Genkin and Engel, 2020). Furthermore, explicitly capturing non-stationary stochastic dynamics is essential to fully uncover biological computations (Genkin et al., 2021). Our DMM addresses these challenges: it shares the goal of trajectory inference but differs in a critical respect by explicitly parameterizing a stochastic transition model with a learned noise structure, rather than inferring a deterministic dynamical system with additive noise. This distinction is not merely technical—it is what enables the model’s stochastic dynamics to generate the full distribution of reaction times, rather than merely smoothing the observed trajectories. The noise is not a nuisance to be filtered; it is the source of behavioural variability. Looking forward, replacing the LSTM-based inference network with a Transformer architecture (Candelori et al., 2025) may further improve long-sequence inference and real-time decoding, though the core generative framework would remain unchanged.

Several important extensions remain. Our findings are grounded in PMd recordings from two macaques performing a single countermanding task. Whether the same 3-dimensional, Markov structure holds across brain regions involved in the stopping network (supplementary motor areas, basal ganglia), across species, and across richer behavioural paradigms will determine the generality of this computational motif. A further open question concerns the spatial reinterpretation of the Pause-then-Cancel framework proposed above. In the standard temporal account, the Pause is stimulus-locked and fires at every SSD; in our spatial interpretation, the Pause effect is a property of a region of the manifold that becomes geometrically inaccessible at long SSDs. Distinguishing these two possibilities through systematic characterization of the latent force field under different contexts is a natural application of the present framework. The framework itself — using functional completeness to identify the stiff dimensionality of a neural system — is fully general; the specific answer (D = 3, Markov) may be task- and region-dependent.

The Independent Race Model succeeded for decades because its behavioural predictions were approximately correct—independent processes appear to account for the observed patterns of reaction times and stopping probabilities. Our results reveal why: a single, 3-dimensional dynamical engine, constrained by the geometry of the premotor cortex, generates all the behavioural complexity that was previously attributed to separate, independent modules. The apparent independence of Go and Stop was not a property of the underlying computation but an artifact of observing the output without seeing the manifold that produced it. By making that manifold visible, generative, and manipulable, we complement an abstract cognitive-level assumption with an explicit physical mechanism—demonstrating that the gap between neural implementation and behavioural theory can be closed not by building up from single neurons, but by letting the data reveal the low-dimensional algorithm that was always there.

## 4 Methods

### 4.1 Experimental Data and Preprocessing

We analysed multi-unit activity (MUA) from a countermanding reaching task recorded in the left PMd of two macaques. The recordings analysed here are those reported in Pani et al. (2022), which used spike-based measures. The recordings were obtained from two males (Monkey P and Monkey C) rhesus macaque monkey (Macaca mulatta) trained to perform a countermanding reaching task (Logan et al., 1984). Each trial began with the appearance of a central red target. After a variable hold period (400–900 ms), a peripheral target appeared either to the left or right, while the central target disappeared—constituting the Go cue. The monkey was required to initiate a reach toward the peripheral target and maintain contact for reward. The reaction time (RT) was defined as the latency between the Go cue and movement onset.

On Stop trials (20%-30% of the trials), the central target reappeared after a variable stop-signal delay (SSD), instructing the monkey to cancel the ongoing movement. Stop trials were labelled as successful stop when the reach was successfully inhibited after the Go cue, and unsuccessful stop otherwise. SSD was updated by a staircase procedure targeting approximately 50% successful stopping. The realised success rate among stop trials was close to this target for Monkey P. for Monkey C, the analysed sessions yielded a higher rate (67%; see below): one of the sessions (27/05/2014) probed a Go/No-Go-like regime with a predominance of short SSDs, yielding 92% successful and 8% unsuccessful stops, which raises the pooled rate above the ∼50% observed in the remaining sessions. We retained this session rather than excluding it: far from compromising the analysis, it probes the model’s behaviour in a limit condition — where stopping is easy and the Stop process rarely fails — which is itself informative about how the model’s stopping mechanism operates across the full range of task difficulty. Because all analyses are conditioned on trial outcome rather than on the aggregate stopping rate, this heterogeneity in session-level success rates does not affect the results.

For each trial, we consider a 1280 ms window (from 280 ms before the Go cue to 1000 ms after it) of the neural activity recorded from the left dorsal premotor cortex (PMd) using a chronically implanted 96-channel Utah array. Raw signals, sampled at 24.4 kHz, were processed to extract a 96-dimensional multi-unit activity (MUA) time series (band-pass filtering 0.3–7 kHz, full-wave rectification, low-pass smoothing).

For Monkey P, we analyzed three recording sessions (02/12/2013; 09/01/2014; 16/01/2014) occurring within a 45-day window to ensure stability of the recorded neural population. These sessions exhibited comparable RT distributions, indicating a consistent movement-preparation dynamics and permitting data pooling without introducing session-specific behavioral variability. The selected sessions, drawn from a broader 9-month experimental period, comprised a total of 2,693 trials. Likewise, for Monkey C, we included three sessions (24/04/2014; 27/05/2014; 28/05/2014) spanning 35 days and displaying similar RT distributions, chosen from the eight sessions available.

All trials from these sessions were retained, without further exclusion, to preserve the full distribution of behavioural and neural variability. Trials were aligned to the Go cue. No temporal windowing or truncation was applied. The MUA signal (derived as described in Sec. 5.1) was binned at 5*ms* resolution, yielding sequences of 256 time steps per trial. For the purposes of state-space modelling, we further downsampled by retaining one sample every 10*ms*, to reduce high-frequency noise, resulting in sequences of 128 time steps.

Each session was independently normalized: for each of the 96 MUA channels, we applied z-scoring across all trials and time steps within that session. This preserves session-specific baseline activity while ensuring comparability across channels. No smoothing or averaging was applied.

Trials were unevenly distributed in the dataset, with 80% (70%) no-stop trials, 10% (10%) unsuccessful stop trials, and 10% (20%) successful stop trials, and dataset was partitioned into 1,881 (2430) trials for training, 400 (511) for validation, and 412 (547) for held-out testing for Monkey P (Monkey C).

All analyses are performed on a per-animal basis: the selected sessions for each subject are pooled into a single dataset, and the DMM and all readouts are trained and evaluated separately for Monkey P and Monkey C.

The task structure is represented by a 4D one-hot vector corresponding respectively to Go-Left ((1, 0, 0, 0)), Go-Right ((0, 1, 0, 0)), pre-Go ((0, 0, 1, 0)), and post-Stop context ((0, 0, 0, 1)). This design provides the model with the task information required for learning context-dependent dynamics.

### 4.2 Generative Dynamical Model: Deep Markov Model (DMM)

We modelled the neural population dynamics using a Deep Markov Model (Krishnan et al., 2017). In this framework, the observed variable *x*_*t*_ ∈ ℝ^96^ denotes the normalized multi-unit activity (MUA) at time *t*, while *z*_*t*_ ∈ ℝ^*D*^ represents a low-dimensional latent state capturing the underlying population dynamics, with *D* ∈ {2, 3, 4} (see Results). Task context is encoded by a one-hot vector *c*_*t*_, defined at the end of the previous subsection, which specifies which of the four task conditions is active at each time point. According to this formulation, we define a latent conditional Markov process (Norris, 1998) with transition *p*(*z*_*t*_ | *z*_*t−*1_, *c*_*t*_) and emission factors *p*(*x*_*t*_ | *z*_*t*_) parameterized by deep neural networks. For inference, rather than adopting the mathematically principled, non-causal Deep Kalman Smoother *q*(*z*_*t*_ | *z*_*t−*1_, *x*_*t*:*T*_ , *c*_*t*:*T*_), we used the structured (i.e. autoregressive, in contrasts with mean-field approximations that assume full independence), causal inference network of the form *q*(*z*_*t*_ | *z*_*t−*1_, *x*_1:*t*_, *c*_1:*t*_), corresponding to the ST-L inference network in Krishnan et al. (2017), that we refer to as Deep Kalman Filter. This allows genuine online predictions, where the model can infer the latent state up to the current time step and immediately generate its future evolution without contamination from future observations, making it suitable for real time applications and predictive evaluations described in Section 3.

The model has three components (Fig. S1).

1. **Encoder / Inference network**. The encoder allows to probabilistically infer, from a 96-dimensional MUA trajectory, the corresponding low dimensional, latent space trajectory. Implemented as a Deep Kalman Filter, the encoder receives, at each time step, the concatenated neural observation and context [*x*_*t*_, *c*_*t*_], projects them through a linear embedding, and processes the sequence with a one-directional Long Short-Term Memory (LSTM) (Hochreiter and Schmidhuber, 1997). Its output is merged with the previous latent state *z*_*t−*1_ through a Gated Recurrent Unit (GRU) cell (Chung et al., 2014), producing a feature vector used to parameterize the approximate posterior *p*(*z*_*t*_ | *z*_*t−*1_, *x*_1:*t*_, *c*_1:*t*_). We output a full multivariate Gaussian with mean and a Cholesky-factor parameterization of the covariance matrix (diagonal + off-diagonal terms) (Golub and Loan, 2013). Allowing full covariance greatly increases posterior expressivity.
2. **Deep Markov Model (DMM) / Transition model**. This is the simulation core of the model, allowing for the generation of simulated trajectories. *p*(*z*_*t*_ | *z*_*t−*1_, *c*_*t*_) is implemented by conditioning a nonlinear multilayer perceptron (MLP) (Reed and Marks, 1999) applied to *z*_*t−*1_ via a Feature-wise Linear Modulation (FiLM) layer (Perez et al., 2018) generated from the context *c*_*t*_. FiLM modulation allows the context to multiplicatively and additively shift the latent dynamics, leading to more informative and controllable trajectory generation. The transition also outputs a full-covariance Gaussian using a Cholesky factor enforcing a positive-definite covariance. This stochasticity models slow, state-dependent variability in the underlying neural dynamics.
3. **Decoder / Emission model**. This component allows to project the low-dimensional latent states back to the 96-dimensional MUA observation space. The emission model *p*(*x*_*t*_ | *z*_*t*_) is an independent Gaussian decoder: an MLP maps *z*_*t*_ to the mean and log-variance of a diagonal 96-dimensional Gaussian, from which *x*_*t*_ is sampled. This stochasticity accounts for fast, observation-level noise not captured by the latent dynamics, such as residual variability in the MUA signal and measurement noise. The combination of stochastic dynamics and stochastic decoding yields a principled decomposition of neural variability.

### 4.3 Loss Function and Training

The model is trained by minimizing a parametrized extension of the negative Evidence Lower Bound (ELBO) loss (Krishnan et al., 2017), which decomposes into a reconstruction term and a regularization term weighted by a *β* parameter:

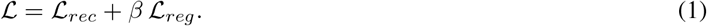

#### Reconstruction term

For a Gaussian decoder with diagonal covariance, the reconstruction loss is the following Gaussian negative log-likelihood:

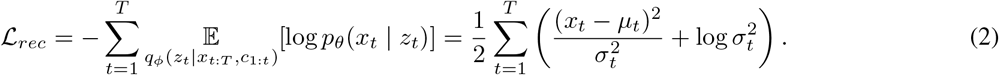

#### Regularization term

The regularization term, in our case, results to be:

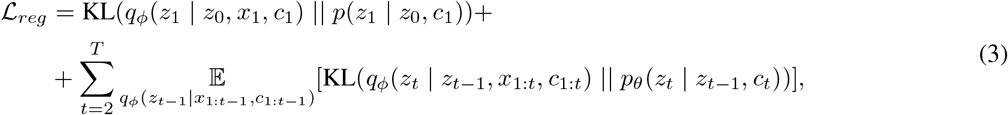

where *T* = 128, *z*_0_ is set to the null vector (*e*.*g*., in 3D, *z*_0_ = (0, 0, 0)), and for two multivariate Gaussians with means *µ*_*q*_ and *µ*_*p*_ and covariances Σ_*q*_ and Σ_*p*_, the KL divergence is (Murphy, 2012):

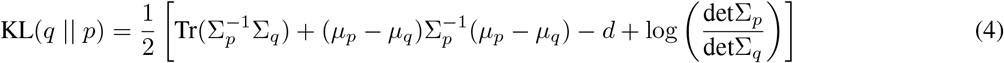

#### Training details

The model was trained for 150 epochs (each epoch corresponding to a number of sampled batches equal in size to the full training set) with batch size 32 for Monkey P (64 for Monkey C), using AdamW (Loshchilov and Hutter, 2017) with weight decay 10^*−*3^ and One-Cycle LR scheduling (Smith and Topin, 2019) with maximum learning rate 10^*−*3^. The dataset was split into 70% training, 15% validation, and 15% testing. To mitigate class imbalance during training, we sampled equal proportions of each trial type in every batch, thereby preventing dominance by the majority class. Crucially, behavioural variables such as reaction times were never used in any part of the loss function or model input. The DMM is trained exclusively on neural activity and task-event context.

#### *β*-DMM weighting

We annealed the beta coefficient over the first 40 epochs, following common KL-warmup strategies for VAE/DMM models (Sønderby et al., 2016; Bowman et al., 2016) to avoid posterior collapse (He et al., 2019; Lucas et al., 2019). Unlike classical *β*-VAE formulations (Burgess et al., 2018), where *β >* 1 pushes the posterior toward a fixed isotropic prior to promote disentanglement, in our model the prior is learned: the transition distribution *p*(*z*_*t*_ | *z*_*t−*1_, *c*_*t*_) is non-isotropic and jointly optimized with the encoder. Thus, increasing *β* pushes the model to rely more on the dynamical predictability embedded in the latent states and less on the information conveyed by observations. This leads to a more self-contained latent dynamics, allowing for long-horizon generative rollouts, though at a cost in reconstruction accuracy. The logic of this selection deserves emphasis, as it constitutes a principled model-selection criterion rather than a post-hoc parameter sweep. In our DMM, reconstruction accuracy is largely insensitive to *β* across a broad range (*β* ∈ [1, 10]): the decoder can compensate for increased regularization without substantial loss in descriptive fidelity. However, *β* directly controls the dynamical self-consistency of the latent process: higher *β* forces the model to rely more on its learned transition rules and less on instantaneous observations, producing latent dynamics that are more autonomous and thus more suitable for generative rollout. We therefore selected the largest *β* for which test-set reconstruction MSE remained strictly lower (by ∼ 1-2%) than the baseline obtained with a 3D PCA representation (specifically, *β* = 4 for both monkeys). This criterion maximizes the model’s capacity for self-driven temporal evolution — the very property needed for generative simulation — without sacrificing descriptive performance.

### 4.4 Comparative and Readout Models

To assess how effectively the DMM latent space capture task-relevant information and the temporal structure of neural dynamics, we trained a set of predictive and functional readout models directly on the inferred trajectories *z*_1:*T*_ . These analyses (i) quantify how reliably behavioural variables can be decoded from the latent space and (ii) assess the dynamical predictability of the latent process itself. All evaluations were performed on held-out test data.

As a baseline comparison, we performed identical analyses on latent representations obtained via Principal Component Analysis (PCA) (Jolliffe, 2002), a standard linear dimensionality-reduction method widely used in neural data analysis (Kobak et al., 2016). PCA was fitted on the DMM training data, reshaped into a two-dimensional matrix (trials × time steps, channels), and the learned linear projection was applied to the test set. This procedure ensures that DMM and PCA latents were compared under identical conditions and matched dimensionality.

#### Functional readout models

These readouts operate hierarchically rather than redundantly. On any trajectory, the movement detector first classifies whether a movement is executed; only if it is does the reaction-time detector regress the corresponding RT (both architectures detailed below). This cascade (classify, then regress conditional on movement) is cleaner and better-posed than a single regressor forced to emit an undefined value on inhibited trials, where no reaction time exists.

We first trained three LSTM-based readout models designed to evaluate how well different aspects of behaviour can be decoded from the latent trajectories. In addition to serving as diagnostic tools, these readouts are later used to analyse emergent properties of the DMM generative model.

#### Reaction time (RT) detector

An LSTM-based regressor predicts a continuous, scaled reaction time from the full latent trajectory *z*_1:*T*_ of no-stop trials. The model consists of a bidirectional LSTM backbone followed by a two-layer regression head and is optimized using MSE loss.

#### Movement detector

A binary LSTM classifier predicts whether a Stop signal results in movement or inhibition, given the full latent sequence *z*_1:*T*_ . The architecture mirrors that of the RT regressor but terminates in a classification head producing a move probability *p*(move), and is trained using binary cross-entropy loss.

#### Direction detector

A binary LSTM classifier predicts movement direction (left vs. right) in no-stop trials from the full latent trajectory *z*_1:*T*_ . The model shares the same backbone structure as the movement detector and outputs a rightward movement probability *p*(right), trained using binary cross-entropy.

#### Latent dynamical predictability

In the characterization of a dynamical model it is paramount to ask how much information about the future state of the system is embedded in its history. To probe this aspect, we trained two classes of latent-space predictors on both DMM and PCA representations.

#### Markov predictor

A two-layer (MLP) mapping [*z*_*t−*1_, *c*_*t*_] to *z*_*t*_, isolating strictly first-order transition structure.

#### Non-Markov predictor

A single-layer LSTM receiving the full past sequence [*z*_1:*t−*1_, *c*_1:*t*_], providing an operational upper bound on predictability by leveraging long-range temporal information.

Both predictors were trained on latent trajectories from the training set and evaluated on held-out test trajectories.

To compare prediction performance across latent spaces with potentially different scales and dimension *D*, we evaluated predictors using an innovation-normalized metric. Defining the latent innovation as Δ*z*_*t*_ = *z*_*t*+1_ −*z*_*t*_ and its predicted equivalent as 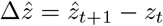 (both assumed to have dimension *D*), we computed the mean squared prediction error in a whitened innovation space:

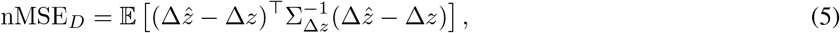

which is equivalent to the average squared Mahalanobis distance (Johnson and Wichern, 2007) between predicted 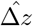 and true Δ*z* innovations.

For interpretability, we report the corresponding explained dynamical variance

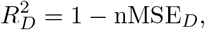

which quantifies the fraction of innovation variance explained relative to a persistence baseline 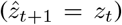. This normalization renders prediction errors invariant to linear reparameterizations of the latent space, enabling fair comparisons between DMM and PCA representations with matched dimensionality but different coordinate systems.

### 4.5 Analysis and Simulation Procedures

To determine the minimal latent dimensionality required for functional completeness of the DMM, we trained models with latent dimensionality *D* ∈ {2, 3, 4} and evaluated them using the full set of behavioural and dynamical metrics described in Section 4.4. The hyperparameter *β* was selected for *D* = 2 and *D* = 4 in order to closely match the reconstruction MSE of the model with *D* = 3. This procedure was devised to isolate how latent dimensionality contributes to the encoding of dynamical information, keeping constant the reconstruction capacity.

In addition to behavioural decoding and latent dynamical predictability, we performed a variance analysis of the inferred latent trajectories. Specifically, we applied Principal Component Analysis (PCA) to the latent trajectories inferred by the *D* = 4 DMM encoder on the training set. The resulting linear transformation was then applied to the inferred latent trajectories of the test set, and the explained variance ratio of each principal component was computed, to give a measure of how dispersed the trajectories are in different directions in the 4D space.

We further tested the generative capabilities of the DMM.

#### Generative reproduction of behavioural variability

We generated synthetic latent trajectories by rolling out the Deep Markov Model *p*(*z*_*t*_ | *z*_*t−*1_, *c*_*t*_). Each rollout was initialized from the inferred latent state at the Go cue of held-out test trials. The generated trajectories were analysed in two complementary ways. First, they were passed through the trained RT detector (Section 4.4) to obtain a synthetic reaction time distribution, which was compared to the empirical distribution using the Kolmogorov–Smirnov test (Wasserman, 2006). Second, multi-unit activity was reconstructed from the generated latent trajectories and compared to the empirical MUA from test trials by examining the mean activity (averaged over channels) across time (from Go cue to 1000*ms* thereafter) and across trials ordered by increasing reaction time.

#### Predictive consistency of generated trajectories

To further assess whether the DMM-generated dynamics preserve trial-by-trial behavioural structure, we evaluated the ability of partially generated latent trajectories to predict reaction times at different stages of the trial. Specifically, we simulated latent trajectories starting from progressively later time points spanning from before to after the Go cue. For each simulation start time *t*_*s*_ ∈ {−280*ms*, … , 1000*ms*} relative to the GO cue (intervalled by 10 ms, with negative values indicating start times preceding the Go cue), we selected only test trials whose true reaction time occurred after *t*_*s*_. Simulation was discontinued once fewer than 120 trials satisfied this criterion.

For each eligible trial and *t*_*s*_, we inferred the corresponding latent state via the DMM encoder and generated 20 independent latent rollouts using the learned transition model, starting from the latent state at *t*_*s*_ and continuing until the end of the trial. The 20 realizations were then averaged to obtain a single mean latent trajectory per trial, reducing stochastic variability induced by sampling. Reaction times were subsequently predicted from these generated trajectories using the trained RT detector (Sec. 4.4). A small fraction of trials yielding undefined predictions (NaNs), zero reaction times, or reaction times coinciding with the simulation start was excluded from further analysis.

For a given *t*_*s*_, we computed the Pearson correlation coefficient between the predicted and true reaction times across trials. This procedure yields a time-resolved measure of how well the DMM-generated dynamics, conditioned on progressively longer segments of the inferred latent trajectory, retain predictive information about behavioural outcomes.

### 4.6 Stop Trials and Virtual Stop Interventions

We next incorporate stop trials to examine how the DMM represents inhibitory control. Stop trials were divided into two classes based on behavioural outcome. Successful stop trials are those in which the Stop signal is effective in inhibiting the planned movement. In contrast, unsuccessful stop trials occur when the movement is executed despite of the Stop signal.

Typically, successful stop trials are associated with Stop signals delivered early after the Go cue; an unsuccessful trial, on the other hand, typically occurs with a late Stop signal, when the movement process has already progressed beyond a point of no return and execution cannot be prevented.

To characterize the latent dynamics underlying stopping, we analyzed both single-trial trajectories and population-level structure at the time of the Stop signal. For visualization, we selected representative successful and unsuccessful stop trials and plotted their inferred latent trajectories in latent space. In addition, we collected the latent states at the Stop time from all stop trials in the training set and examined their distribution in latent space. We performed a linear discriminant analysis in order to derive a linear boundary that maximally separates the two classes in latent space, calculating its accuracy using stops of the test set. For visualization purposes, all latent-space plots show the projection onto the first two latent components. Among the three latent dimensions, the second component carries the strongest discriminative information for both movement direction (left/right) and stopping outcome (successful stop vs. no-stop).

To showcase the capability of the model to simulate ‘what-if’ scenarios, we performed in silico perturbation experiments in which no-stop trials were converted into stop trials. For a given no-stop trial, we inferred the latent trajectory, via the DMM encoder, up to an arbitrary selected virtual Stop time. Starting from the inferred latent state at that time, we switched the context to Stop and generated the subsequent latent trajectory using the DMM *p*(*z*_*t*_ | *z*_*t−*1_, *c*_*t*_). The generated trajectories were then passed to the movement detector of Sec. 4.4 to classify the resulting behaviour as movement or inhibition.

### 4.7 Simulated Stop-Failure Protocol and Context Independence Test

To test context independence—the assumption that the Go reaction-time distribution is identical on stop and no-stop trials—we devised a simulated stop-failure protocol that uses the generative engine to predict the reaction time a Go trial would have produced under a failed stop. For each no-stop trial *i* in the held-out test set, with behavioural reaction time RT_*i*_ and neural trajectory 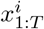, we inferred the latent trajectory 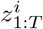 via the DMM encoder. For a given stop-signal delay SSD, we set the latent state at 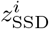 (the inferred state at the time corresponding to SSD after the Go cue), switched the context vector to Stop, and generated *K* = 200 independent forward simulations of the latent dynamics using the DMM transition model *p*(*z*_*t*_ | *z*_*t−*1_, *c*_*t*_).

Each of the *K* simulated trajectories was classified by the movement detector (see Sec. 4.4) as a successful stop (*p*(move) *<* 0.5) or a failed stop (*p*(move) ≥ 0.5). For each failed-stop simulation *k*, the full latent trajectory—composed of the inferred segment 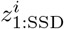 concatenated with the generated segment—was passed through the RT regressor (Sec. 4.4) 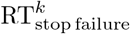 to obtain a simulated stop-failure reaction time RT^*k*^ . For trial *i* at the given SSD, we then averaged over all failed-stop realizations:

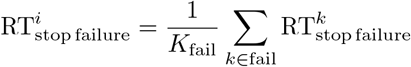

where *K*_fail_ is the number of simulations classified as failed stops. To ensure more reliable estimation, trials for which fewer than 5% of simulations (fewer than 10 out of 200) resulted in a failed stop at a given SSD were excluded from the analysis at that SSD. A trial excluded at one SSD could be retained at another.

For each surviving trial *i* at a given SSD, we computed the per-trial deviation:

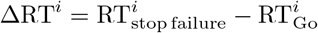

where 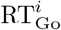 is the trial’s own behavioural Go reaction time. The mean ΔRT across all surviving trials at a given SSD, together with its standard error, quantifies the degree to which the stop process distorts the Go RT distribution at that delay. Under context independence, ΔRT should not be statistically distinguishable from zero at any SSD.

This procedure was repeated for each SSD in the set used during the experimental staircase procedure (six values for each monkey). For Monkey P, the shortest SSD was excluded from the final analysis because only three test trials produced a stop-failure rate exceeding 5% at that delay.

### 4.8 Stochastic Independence Test and SSRT Estimation

We next asked whether the model predicts a dependence of the stop-signal reaction time (SSRT) on the intrinsic speed of the ongoing movement process (as determined by the RT). To this end, we used the generative engine to simulate virtual stopping at a dense range of stop times relative to movement onset, and derived a per-trial estimate of SSRT as follows. Unlike the integration method, which assumes a fixed SSD and returns *RT* ^∗^ as a quantile of the aggregate Go RT distribution, the inverse problem was considered here: fixing each trial’s own RT and asking what SSD would have been “one SSRT away” from it. Mechanistically, SSRT is the time the stop process takes to complete inhibition after the stop signal; if the stop signal arrives exactly one SSRT before the RT, the Go and Stop processes finish at nearly the same moment, placing the trial at the tipping point between successful and unsuccessful inhibition (*p*_Move_ = 0.5). Accordingly, the critical SSD associated with this tipping point was estimated.

For each no-stop trial of the held-out test-set, the latent trajectory was inferred via the DMM encoder multiple times (*n* = 20) and averaged to obtain a mean latent trajectory 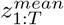. Next, we defined a set of virtual stop times relative to movement onset (RT),

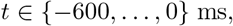

corresponding to 60 discrete time steps with a temporal resolution of 10 ms. For each virtual stop time *t*, we initialized the latent state at 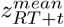 and generated the remainder of the trajectory using the Stop context, multiple times (*n* = 20). Each generated latent trajectory was then passed to the movement detector described in Sec. 4.4, which returned the probability of movement execution, *p*_Move_. For each trial and each virtual stop time, *p*_Move_ was averaged across 20 simulations, yielding a *p*_Move_ curve that characterizes the movement probability as a function of stop timing at the single-trial level.

For each trial, as discussed above, we defined the *critical SSD* as the latest virtual stop time at which the single-trial *p*_Move_ curve remains below 0.5. The per-trial SSRT then follows directly as SSRT = RT − SSD_critical_. Because SSRT is defined relative to RT, sweeping the stop time is naturally expressed as a sweep backward from movement onset. Trials for which this quantity could not be reliably estimated (NaNs) were excluded from subsequent analyses.

To avoid edge effects, i.e. atypical trials with very short or very long reaction times that could bias the analysis, we removed the 5% of trials with the shortest and the 5% with the longest true RTs. We excluded a small number of trials (17 out of 293) for which the estimated SSRT was equal to zero.

Note that this procedure estimates the tipping point of the distribution of possible outcomes consistent with the trial’s inferred state at the simulatiom start, rather than directly recovering the finishing time of the specific stop process that would have occurred on that single, noise-realized trial, a quantity that is fundamentally unobservable, since each real trial reflects a single draw from the underlying stochastic dynamics. We mitigate the resulting per-trial noise by aggregating SSRT estimates, sorted by true RT, over nine groups of trials of equal size, rather than interpreting individual-trial values in isolation. For each group, we computed the mean true RT and the mean estimated SSRT, with error bars given by the standard error of the mean. A linear regression was fitted to these points to quantify the relationship between SSRT and RT.

Finally, to estimate the population-average SSRT, we averaged the *p*_Move_ curves across all trials, obtaining the mean probability of movement execution as a function of time before RT. The average SSRT was defined as the latest time point before the average *p*_Move_ curve crosses the 0.5 threshold.

### 4.9 Virtual stimulation of observed neural activity

The DMM can be used to design interventions at the neural activity level aimed to predictively affect behaviour (notably, reaction times). Such interventions take the form of virtual stimulation protocols aimed to nudge the future neural activity on the basis of the past activity, and are thus potentially reproducible in real neurophysiological experiments.

#### Stimulation protocol

The idea behind these interventions is to indirectly ‘push’ the latent trajectory at time *t* in a trial-independent direction Δ*z*_*t*_ associated with longer *RT* s. Such push increases (or decreases, in the opposite direction) the average *RT* . This ‘latent-space stimulus’ Δ*z*_*t*_ is mapped into a trial-specific desired variation of the standardized MUA activity Δ*x*_*t*_ by inverting the approximate (degerate) linear relationship:

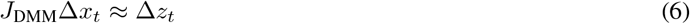

where *J*_DMM_ is the Jacobian of the DMM encoder evaluated at (*x*_1:*t*_, *z*_*t−*1_). This inverse problem is solved using Elastic-Net regression, promoting smaller (in absolute value) and sparser solutions. The Δ*x*_*t*_ actually used in the simulation are determined by rescaling the result of the inversion by a time- and trial-independent factor. Such factor modulates the effect of the stimulation protocol on the *RT* . Given the linearity of Eq. 6, this rescaling is equivalent to rescaling the trial-independent vectors Δ*z*_*t*_.

The protocol prescribes a non-null stimulation only in the time interval *t* ∈ [GO − 100 ms, GO + 90 ms]; outside this range, Δ*z*_*t*_ (and, consequently, Δ*x*_*t*_) is assumed null.

The perturbed input

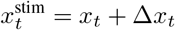

is then concatenated to *x*_1:*t−*1_, alongside *z*_*t−*1_, and encoded to obtain 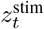. From 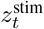 we perform a one-step simulation of the DMM dynamics to get the next hidden state *z*_*t*+1_ and, from that, 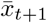 as the mean output of the DMM decoder. 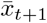 represents the expected neural activity at time *t* + 1 given the history, up to time *t*, of the neural activity subjected to the stimulation protocol. To preserve fast, trial-specific variability that is not captured by the latent states, and thus ensure a meaningful comparison with true neural trajectories, we add to 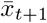 the actual model’s residual *δx*_*t*+1_ for the specific trial at time *t* + 1. In other words, *δx*_*t*+1_ assumes the role of additional noise.

Such residual is the difference between the true *x*_*t*+1_ and the average DMM reconstruction at time *t* + 1. This latter is obtained by inferring from *x*_1:*t*+1_ 20 latent trajectories 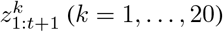; each latent trajectory is decoded into an inferred neural trajectory 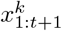; all these neural trajectories are averaged to get the residual:

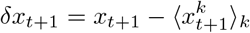

#### Determination of Δ*z*_*t*_

Given a trial *i*, with reaction time *RT*_*i*_ and MUA 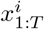, we sample, via the DMM encoder, an inferred latent trajectory 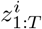. Given a time *t*, we determine direction Δ*z*_*t*_ of maximum *RT* increase by fitting:

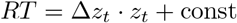

on the 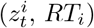 pairs; · represents the scalar product of vectors.

## Acknowledgments

The activity has been funded by the European Union – Next Generation EU under the Italian Ministry of University and Research (MUR) project ECS00000024 “Ecosistemi dell’Innovazione” – Rome Technopole, public call n. 3277, PNRR – Mission 4, Component 2, Investment 1.5.

## Supplementary Material

This section provides additional model details, robustness checks, and control analyses supporting the results presented in the main text. Sec. 1.1 provides an intuitive scheme of the generative and inference structure of the Deep Markov Model. Sec. 1.2 provides a control ruling out over-regularization as the cause of the fourth latent dimension’s negligible contribution. Sec. 1.3 illustrates, at the level of a single trajectory, the in silico intervention protocol underlying the context-independence test. Sec. 1.4 and 1.5 reports control analyses for the virtual stimulation and stochastic independence experiments. Finally, Sec. 1.6

### 1.1 Deep Markov Model: generative and inference structure

Figure S1 illustrates the three components of the Deep Markov Model described in Methods, Sec. 4.2: the stochastic transition model, the causal inference network, and the emission (decoding) model. Together, these define how latent states are generated, inferred, and mapped back to multi-unit activity.

**Fig. S1:**
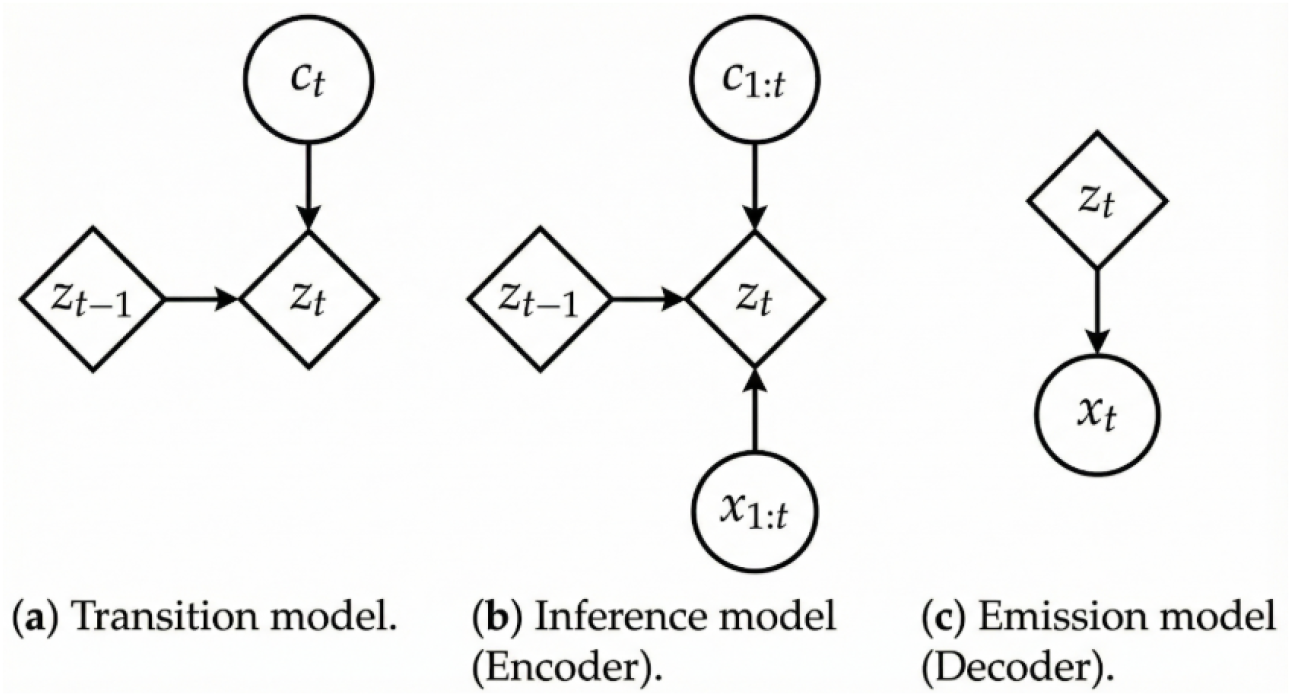
(a) The generated *z*_*t*_ depends on previous *z*_*t−*1_ and current context *c*_*t*_. (b) The inferred *z*_*t*_ depends on the previous *z*_*t−*1_, and on all input observations *x*_1:*t*_ and context *c*_1:*t*_ up to the present. (c) The decoded *x*_*t*_ depends only on *z*_*t*_.

### 1.2 The fourth latent dimension is suppressed by the data, not by regularization

In the main text (Sec. 2.2), we identified *D* = 3 as the minimal dimensionality for functional completeness, noting that a 4th latent dimension contributes negligible explained variance under the DMM trained with *β* = 8. Because *β* controls the strength of the KL regularization term, a natural concern is that this suppression might simply reflect over-regularization (i.e., the prior forcing unused dimensions toward zero) rather than a genuine property of the underlying neural data. To rule this out, we retrained the 4D DMM with a substantially reduced *β* (*β* = 0.5, a 16-fold reduction relative to the main analysis) and repeated the explained-variance analysis.

**Fig. S2:**
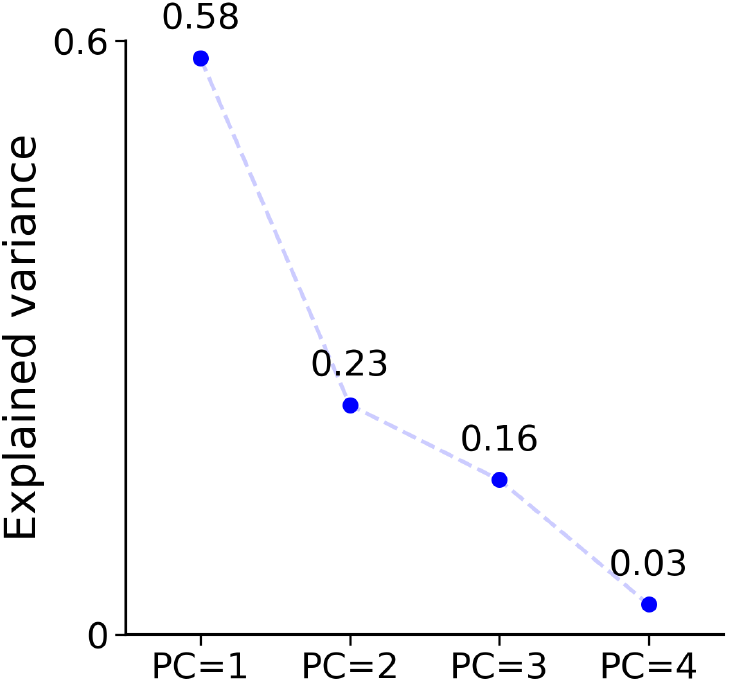
Retraining the 4D DMM with a reduced *β* (*β* = 0.5, compared with *β* = 8 used in the main analysis) and repeating the explained variance analysis. The fourth latent dimension contributes negligibly to the system dynamics, confirming that its suppression is driven by the data rather than by over-regularization.

As shown in Fig. S2, the fourth dimension remains negligible even under this much weaker regularization, confirming that its suppression reflects the intrinsic dimensionality of the underlying neural dynamics rather than an artifact of the *β*-weighted training objective.

### 1.3 In silico conversion of a successful Stop trial into a No-stop trial

To illustrate, at the level of a single trajectory, another possible in silico intervention, in addition to the ones described in Sec. 2.4, Fig. S3 shows an example in which a successful Stop trial is perturbed to ignore the Stop signal, effectively converting it into a simulated No-stop movement trajectory.

**Fig. S3:**
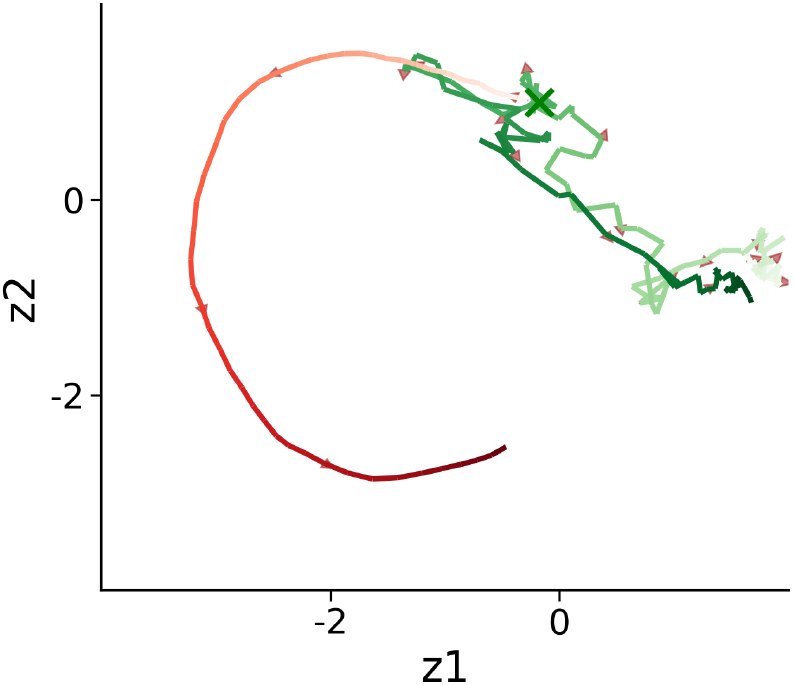
Successful Movement: A successful stop trial trajectory (green) is perturbed to ignore the Stop signal, effectively transforming it into a no-stop movement trajectory (red). From the SSD onward, we continue to provide the model with the contextual signal it received immediately after the GO cue.

### 1.4 Control analyses for the virtual stimulation experiments

The virtual stimulation experiments of Sec. 2.4 (Fig. 9) show that targeted, in silico perturbations of MUA induce systematic, sign-specific shifts in predicted reaction times. Here we report two control analyses ruling out trivial explanations for these shifts: regression to the mean, and stimuli larger than the intrinsic scale of neural variability.

**Fig. S4:**
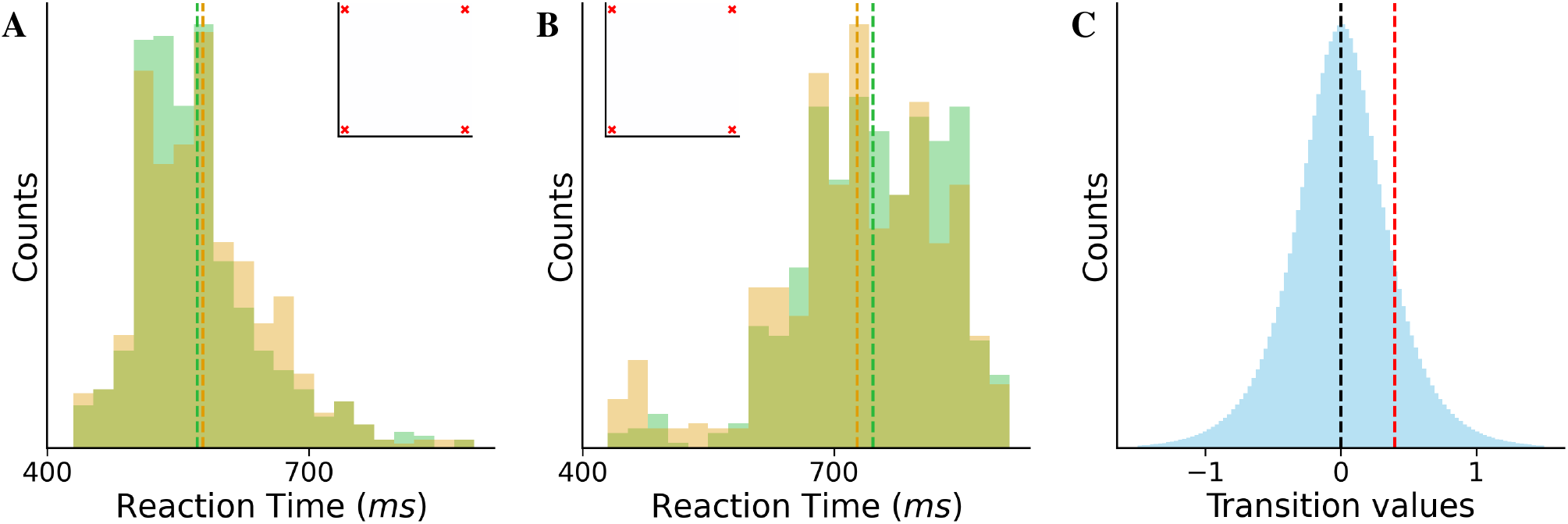
**(a**,**b)** Reaction time (RT) distributions obtained from DMM simulations initialized at *GO* + 100 ms in the absence of stimulation, for trials selected for RT-lengthening (a) and RT-shortening (b). The distribution of predicted RT from infinitesimally perturbed DMM rollouts (amber) match the distribution of RTs predicted from unperturbed DMM rollouts (green). Vertical dashed lines indicate the respective distribution means. *Insets* show the applied input-space perturbation, corresponding to a zero-valued stimulus. **(c)** Distribution of per-step neural activity changes, aggregated across trials, time points, and channels (Δ*x*_*t*_ = *x*_*t*+1_ − *x*_*t*_). The black dashed line marks the mean of the distribution (approximately zero), while the red dashed line indicates the standard deviation (*σ* ≈ 0.4), providing a reference scale for the magnitude of the applied stimuli in the main experiment.

Panels A and B confirm that, absent any stimulation, infinitesimally perturbed rollouts reproduce the unperturbed RT distribution for both the trials selected for RT-lengthening and RT-shortening, ruling out regression to the mean as an explanation for the shifts reported in Fig. 9. Panel C provides a reference scale (*σ* ≈ 0.4) for the intrinsic, per-step variability of the neural activity, against which the magnitude of the applied stimuli in the main experiment can be judged.

### 1.5 Robustness of the RT–SSRT correlation to single-trial noise

A possible concern about the positive correlation between RT and per-trial SSRT reported in the main text (Fig. 8) is that it could arise as a statistical artifact of the estimation procedure itself, rather than reflecting a genuine property of the underlying race dynamics. Because the per-trial SSRT is computed as SSRT = RT − SSD^∗^, any single-trial noise contributing to a trial’s observed RT could, in principle, propagate directly into the SSRT estimate, mechanically inducing a positive correlation between the two quantities independent of any true coupling between the Go and Stop processes.

To test this possibility, we repeated the analysis of Fig. 8 using, in place of each trial’s single observed RT, the simulated RT obtained from the model’s own generative dynamics: starting from each trial’s inferred latent state at its estimated SSD^∗^, we generated 30 independent stochastic rollouts under the Go context, passed each through the RT detector (Sec. 4.4), and took the mean of the resulting 30 RTs as the trial’s simulated RT. Because this quantity is averaged over many independent stochastic realizations from the same latent state, it is largely decoupled from the specific noise realization of the single real trial, and any correlation with SSRT emerging from this analysis cannot be explained by direct propagation of single-trial noise through the RT − SSD^∗^ formula.

Figure S5 shows the resulting SSRT-vs-simulated-RT relationship for Monkey P. The correlation closely reproduces both the trend and the approximate range obtained using the true, single-trial RTs, with a regression slope clearly different from the value of 1 expected under pure algebraic noise propagation. This indicates that the RT–SSRT relationship reported in the main text is not primarily driven by single-trial noise leaking into the SSRT estimate, and supports interpreting it as reflecting a genuine property of the model’s latent race dynamics rather than an artifact of the estimation procedure.

**Fig. S5:**
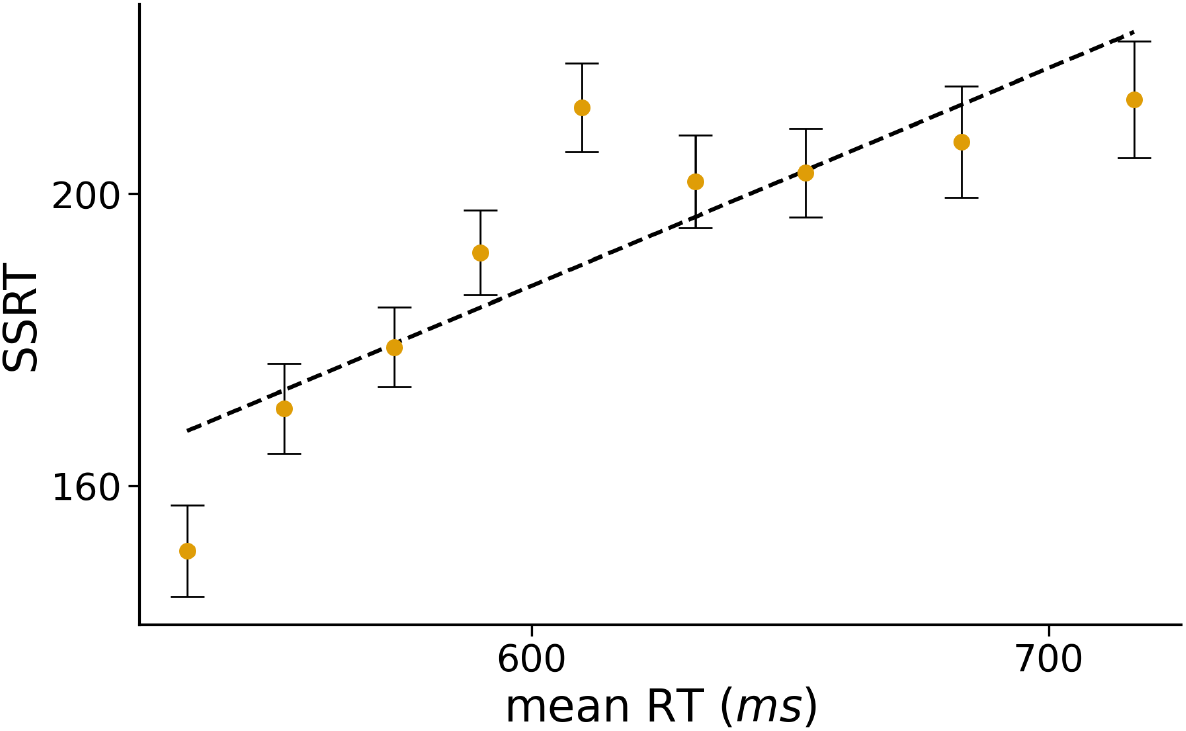
**The RT–SSRT correlation is robust to single-trial noise**. SSRT plotted against the trial-averaged simulated RT (mean of 30 stochastic rollouts per trial, generated from each trial’s inferred latent state at its estimated SSD^∗^; see Sec. 1.5), for Monkey P. Error bars denote standard error within each RT bin. The dashed line shows the linear regression fit. The relationship closely matches that obtained using true, single-trial RTs (Fig. 8), with a slope distinct from 1, indicating that the correlation is not attributable to single-trial noise propagating algebraically from RT into the SSRT estimate.

### 1.6 SSD-dependent SSRT: a signature of context independence violation

We further test context independence investigating the SSRT-vs-SSD profile (Fig. S6), which provides the most immediately interpretable signature for an experimentalist—and a direct demonstration that the integration-method SSRT depends strongly on the chosen SSD.

We performed a fully *in-silico* experiment. We first generated, supported by the result in Fig. 4A, a number of no-stop RTs – much larger than what typically can be obtained in single-subject experiments – as a reference empirical distribution to use with the integration method to estimate SSRT. We then ran, for each SSD in the range [215, 490] ms, in steps of 25 ms, the stop-simulation protocol described in Sec. 1.7 and recorded the fraction *q* of simulated trajectories that resulted in a movement. Such SSD range was chosen as being common to both animals, allowing a direct comparison between the two, without extrapolating beyond experimentally sampled values. Four reference values of *q*, approximately 0.10, 0.30, 0.60, and 0.80, corresponding to inhibitory probabilities of 90%, 70%, 40%, and 20%, respectively, were then used to select four representative SSDs (SSD_1_ through SSD_4_) in direct analogy with the individually tailored SSD schedule used by Gulberti et al. (2014). Matching on inhibition probability rather than on raw SSD is essential for a fair comparison: because the inhibition function depends on each individual’s own RT distribution, a given SSD corresponds to different inhibitory difficulty across subjects and species. Anchoring to fixed inhibition probabilities (90/70/40/20%) places all subjects on a common functional axis, enabling a genuine cross-species comparison. For each of these four SSDs the SSRT was estimated as the *q*-th quantile of the simulated no-stop RT distribution minus the SSD (see Sec. 1.7). This whole pipeline was repeated independently for the two monkeys.

**Fig. S6:**
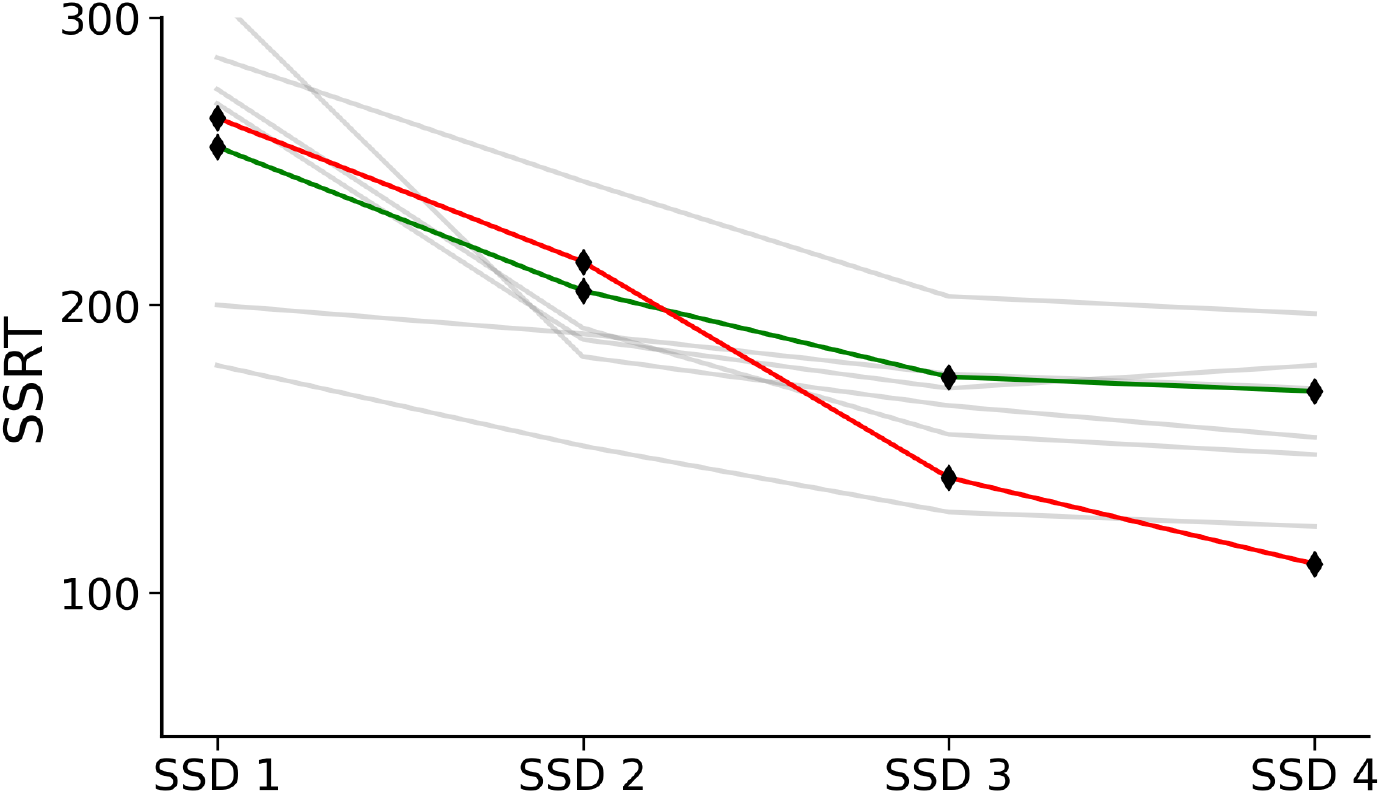
**Model-derived SSRT as a function of SSD matches human behavioural benchmarks**. Stop-Signal Reaction Time (SSRT, in ms) plotted against four reference Stop-Signal Delays (SSD_1_–SSD_4_), defined as the SSD at which the fraction of simulated movements equals *q* ≈ 0.10, 0.30, 0.60, and 0.80, corresponding to inhibitory probabilities of 90%, 70%, 40%, and 20%, respectively. Specifically, they resulted in: SSD_1_ = 265 ms (215 ms), SSD_2_ = 365 ms (265 ms), SSD_3_ = 465 ms (440 ms), SSD_4_ = 490 ms (490 ms) for Monkey P (Monkey C), depicted in the figure as the green (red) curve. Grey lines: SSRT values for six human participants from Table 6 (Manual / Hands condition) of Gulberti et al. (2014), at the individually calibrated SSDs yielding 90%, 70%, 40%, and 20% inhibition probability. Both animals reproduce the monotonically decreasing SSRT-vs-SSD trend observed in humans, and their values remain within the envelope of the human data across all four reference delays. Comparability across species rests on matching inhibition probability, not raw SSD, and is intended as a qualitative check of a well-established regularity, not a claim of quantitative cross-species equivalence. The SSRT was computed as *q*^th^ quantile of the simulated no-stop RT distribution minus the SSD (see Sec. 1.7).

The resulting SSRT-vs-SSD profiles for both Monkey P and Monkey C are shown in Fig. S6, together with the values reported for six human participants in Table 6 of Gulberti et al. (2014) (Manual / Hands condition), which span the range [≈ 120, 310] ms and all exhibit a monotonically decreasing trend with SSD. Both simulated animals reproduce the same qualitative pattern: SSRT decreases monotonically as the SSD increases, in line with what is expected when higher SSDs are paired with higher inhibitory difficulty. This finding is fully consistent with the SSD-dependent SSRT profiles documented in human participants performing hand-movement stop-signal tasks (Gulberti et al., 2014), and with the analogous decreasing trend visible in the supplementary material of Bissett et al. (2021a) (their Fig. S5). The model’s SSRT values at SSD_1_ lie in the upper part of the human range (≈ 255–260 ms), while those at SSD_4_ approach the lower boundary (≈ 115–120 ms), remaining within the envelope defined by the six human subjects throughout.

These results provide an additional and independent, behavioural-level validation of the generative engine. The model was trained exclusively on multi-unit activity and movement direction context—no behavioural quantity related to stopping latency entered either the training objective or the SSRT estimation procedure—yet it spontaneously recapitulates not only the range spanned by SSRT, but also the systematic, SSD-dependent decrease that has been observed across effectors and species. This demonstrates that the model has genuinely internalised the hidden temporal mechanism underlying inhibitory control, rather than merely fitting surface-level statistics of the neural data.

### 1.7 Population-level SSRT estimation at varying SSDs

To characterise the population-level SSRT as a function of SSD we proceeded in two stages.

#### Generating the no-stop RT reference distribution

For each test-set trial, the encoder of the DMM was run 20 times to obtain an ensemble of stochastic latent trajectories up to the Go cue (*z*_GO_). From each *z*_GO_, the generative model was then unrolled forward under the movement-direction context (Left or Right, matching the original trial), provided at its true experimental onset; the recurrent dynamics allow this input to affect the latent state only after the delay expected from visual processing and afferent transmission to PMd, rather than instantaneously. After repeating this procedure for all trials, the resulting trajectories were decoded by the RT detector to produce a distribution of predicted reaction times (the simulated no-stop RT distribution). This distribution was verified to closely match the empirical RT distribution before proceeding (see Sec. 2.3).

#### Sweeping over SSDs

For each candidate *SSD* ∈ {215*ms*, 240*ms*, … , 490*ms*} (in steps of 25 ms), the following procedure was applied to the same ensemble of inferred latent states. First, the generative model was unrolled from *z*_GO_ under the appropriate movement-direction context, until reaching *z*_SSD_. The dynamics were then continued from *z*_SSD_ under the Stop context for the remaining steps of the trial. The resulting trajectories were passed through the movement detector, and the fraction *q* of trajectories classified as a movement was recorded. This fraction estimates the probability of stop failure at that SSD. The SSD range was chosen as the overlap between the experimental SSD distributions of the two animals, ensuring an identical analysis for both subjects.

#### SSRT computation and comparison with human data

The SSRT at each SSD was estimated as

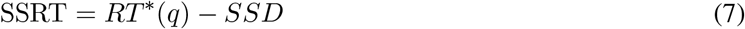

where *RT* ^∗^(*q*) denotes the *q*-th quantile of the simulated no-stop RT distribution. This is equivalent to the integration method of Logan et al. (1984) and Verbruggen et al. (2019), applied at a continuously varying SSD rather than at a single staircase-determined value.

Four reference points—*q* ≈ 0.10, 0.30, 0.60, and 0.80—were selected to identify the SSDs (SSD_1_ through SSD_4_) associated with inhibitory probabilities of 90%, 70%, 40%,and 20%, respectively. This directly mirrors the design of Gulberti et al. (2014), who individually calibrated four SSDs to those exact inhibitory probabilities for each of six human participants (CS, DS, EH, IM, IW, SW) and reported the corresponding SSRTs in their Table 6 (Manual / Hands condition). The resulting SSRT values were then plotted together with the six human profiles to yield Fig. S6.

